# Spatial profiling identifies a GLIS2-associated fibroblast state linked to peritoneal recurrence in gastric cancer

**DOI:** 10.64898/2026.05.21.726806

**Authors:** Seungho Lee, Soohyuk Cho, Dong-Seok Han, Jungmin Kim, Hoon Hur, Hyongbum Henry Kim, Jae-Ho Cheong, Tae-Min Kim

## Abstract

Cancer-associated fibroblasts are transcriptionally heterogeneous, yet how their reprogramming shapes gastric cancer progression remains unclear. Integrating single-cell and spatial transcriptomics, patient-derived fibroblast co-culture, and clinical cohorts, we identified a spatially organized continuum of extracellular matrix-producing fibroblasts. An inferred trajectory connected homeostatic ECMh through inflammatory-chemoattractive ECMi to tissue-remodeling ECMr, with increasing GLIS2 regulon activity and ECMr enrichment near the tumor–stroma interface. ECMr co-culture induced a non-EMT epithelial program characterized by coordinated laminin-332 subunit expression, hemidesmosome-associated genes, a matrix-remodeling protease profile, and reduced proliferative activity. Longitudinal imaging showed progressive cancer-cell elongation and positional displacement over the fibroblast layer. ECMr-associated GLIS2 regulon activity in resected primary tumors was associated with shorter peritoneal metastasis-free survival across two gastric cancer cohorts, with a similar association in colorectal cancer. A peritoneal metastasis recapitulated the corresponding stromal and epithelial organization. Together, these findings define an ECMr-centered stromal–epithelial wound-repair circuit linked to subsequent peritoneal dissemination.

## Introduction

Gastric cancer (GC) remains a major global health concern, ranking fifth worldwide for both cancer incidence and cancer-related mortality^1^. For localized GC, surgical resection remains the principal curative treatment. Although targeted agents and immune checkpoint inhibitors have improved outcomes for selected patients, the overall prognosis of advanced GC remains poor^2,3^. Among the patterns of GC progression, peritoneal metastasis is one of the most lethal, is frequently refractory to systemic therapy, and is associated with a median survival of less than one year^4,5^. The Asian Cancer Research Group (ACRG) molecular classification identified an MSS/EMT subtype associated with the poorest prognosis and frequent peritoneal recurrence^6^. Although this association has commonly been interpreted as reflecting tumor cell-intrinsic EMT, accumulating evidence suggests that bulk EMT signatures can substantially reflect stromal enrichment rather than epithelial-to-mesenchymal conversion within malignant cells^7,8^. This observation redirects attention from tumor cell-autonomous invasion toward stromal-mediated mechanisms; however, the specific stromal programs associated with peritoneal dissemination remain poorly defined.

Cancer-associated fibroblasts (CAFs) constitute a major stromal population in the tumor microenvironment and are particularly abundant in stroma-rich tumors such as diffuse-type gastric cancers. Single-cell transcriptomic studies across multiple cancer types have revealed substantial functional heterogeneity within CAF populations^9–16^. In GC, fibroblast subtypes such as FAP+INHBA+ fibroblasts have been associated with poor prognosis^17^. However, most current classifications define fibroblast subtypes from transcriptomic snapshots and fixed marker sets, providing limited insight into the continuous state-transition dynamics underlying CAF heterogeneity. Fibroblasts are inherently plastic cells capable of dynamically adapting their cellular states in response to microenvironmental cues including tissue injury, inflammation, and malignancy^18^. During physiological wound repair, resident fibroblasts depart from homeostasis and acquire spatially and temporally organized inflammatory and matrix-remodeling state^19,20^. Whether gastric tumors co-opt this reparative plasticity, how the resulting fibroblast states are spatially organized, and how they influence malignant epithelial behavior and peritoneal dissemination remain incompletely understood.

In this study, we integrated publicly available and in-house GC scRNA-seq datasets with spatial transcriptomics, patient-derived fibroblast co-culture, and clinical cohort analyses to resolve the state diversity and spatial organization of ECM-producing fibroblasts. We reconstructed an inferred reprogramming trajectory in which fibroblasts progressed from the homeostatic ECMh state through the inflammatory-chemoattractive ECMi state to the tissue-remodeling ECMr state. We then mapped this continuum along the tumor–stroma axis. We identified ECMr as the remodeling-associated end of the inferred trajectory and showed that co-culture with patient-derived ECMr fibroblasts induced a non-EMT, wound healing-like epithelial program characterized by coordinated expression of laminin-332 (LM332) subunit genes, a hemidesmosome-associated gene program, a matrix-remodeling protease profile, and reduced proliferative activity. This epithelial program was spatially enriched at the tumor–stroma interface, whereas higher ECMr-associated GLIS2 regulon activity in resected primary tumors was associated with subsequent peritoneal recurrence across independent clinical cohorts. Together, our findings define an ECMr-centered stromal–epithelial wound-repair circuit that links fibroblast reprogramming at the tumor interface to a low-proliferative epithelial movement program and subsequent peritoneal dissemination.

## Results

### Fibroblast heterogeneity and spatial organization in gastric cancers

Using-gastric cancer (GC) scRNA-seq data as primary resources (HRA003647, China National Center for Bioinformation), we identified various stromal cell clusters in primary GCs, including six distinct fibroblast subtypes, pericytes (*RGS5, NOTCH3*), and smooth muscle cells (*DES, ACTA2*)^21^ (Figure 1A, Figure S1A and S1B, and Table S2). Among the fibroblast populations, the SOX6+ and APOE+ subtypes were predominantly enriched in normal samples, whereas the MMP1+ subtype was identified as a tumor-specific but also highly sample-specific subtype, *i.e.,* mainly detected in two out of 13 samples (Figure 1B and Figure S1C). In addition, we identified a group of fibroblast clusters characterized by the robust expression of pan-ECM markers, including *OGN*, *DCN*, *LUM*, and *CCDC80* (Figure S1D). Within this group, we resolved three distinct ECM fibroblast cells. These included ECMh (*homeostatic* ECM fibroblast, *CD34*, *CLEC3B*, *MFAP5*), which was more abundant in normal tissues, and two tumor-associated clusters designated as ECMi (*inflammatory*-*chemoattractive*) and ECMr (*tissue remodeling*) (Figure 1C and D). ECMi was characterized by a stress-responsive chemoattractive program marked by *CXCL12*, together with *NAMPT* (NAD salvage/inflammatory stress module) and *ICAM1* (adhesion-related activation), consistent with an early inflammatory, injury-responsive stromal state in tumor tissues (Figure 1D). In contrast, ECMr exhibited relative up-regulation of remodeling-associated markers including *TAGLN*, *DIO2*, and *INHBA* (Figure 1D). To delineate the functional complementarity between these fibroblast clusters, we performed differential pathway activity analysis (Methods). ECMi showed preferential enrichment of pro-inflammatory signaling programs, including TNFα signaling, IL6–JAK–STAT3 signaling, and monocyte chemotaxis (Figure 1E), whereas ECMr was enriched for niche remodeling programs, with high scores for ECM assembly, collagen fibril organization, and angiogenesis (Figure 1E, Table S3). We also validated the robustness of these findings using two independent gastric cancer scRNA-seq cohorts, a publicly available dataset^17^ and an in-house dataset previously generated by our group (Figure S2)^22^. Although the MMP1+ population was not detected, ECMh, ECMi, and ECMr and their defining transcriptional programs were consistently recovered in both validation cohorts. Together, these analyses identify a reproducible set of ECM fibroblast subtypes in GC, comprising a normal-enriched homeostatic state and two tumor-enriched states characterized by inflammatory-chemoattractant and tissue-remodeling programs.

**Figure 1.**
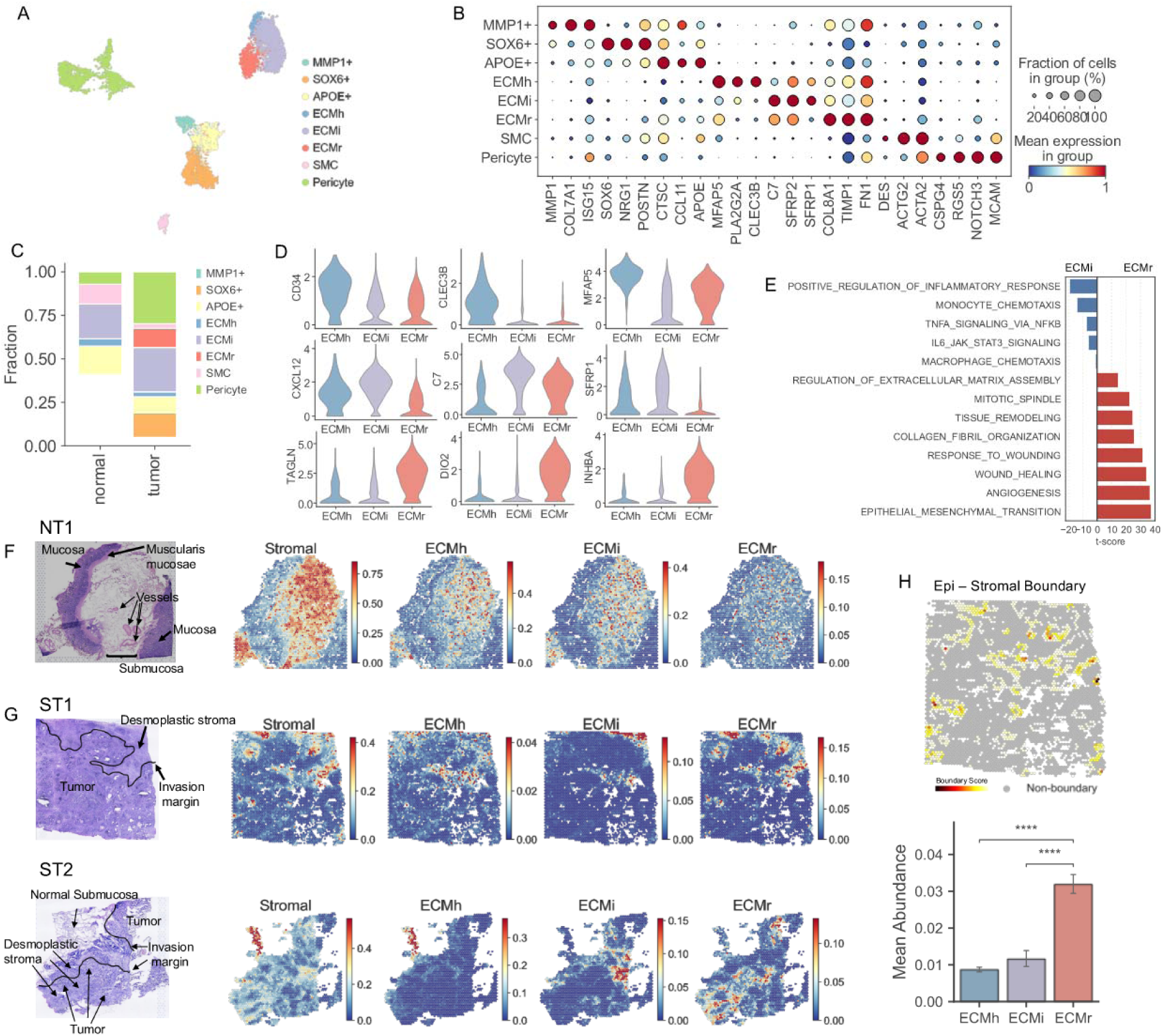
Landscape and Functional Heterogeneity of Fibroblasts in Gastric Cancer. (A) UMAP of stromal cells colored by subset annotation. (B) Dot plot of curated marker gene expression across stromal subsets. Dot size, fraction of cells expressing each gene; color, scaled mean expression. (C) Fractional composition of stromal subsets in normal and tumor tissues, pooled within each condition. (D) Violin plots of subset-defining markers in ECMh, ECMi, and ECMr fibroblasts. (E) Differential pathway activity between ECMi and ECMr fibroblasts. Per-cell scores were inferred with a univariate linear model on curated MSigDB Hallmark and GO Biological Process gene sets, and t-scores were calculated by Welch’s t test (ECMr versus ECMi). (F) H&E image of normal gastric tissue (NT1) and cell2location-inferred spatial maps of the stromal compartment, ECMh, ECMi, and ECMr. Color, estimated cell fraction per spot. (G) H&E images and spatial maps as in (F) for gastric cancer tissues ST1 (top) and ST2 (bottom). (H) (Top) Epithelial-stromal boundary score map in ST1, non-boundary spots in gray. (Bottom) Mean ECMh, ECMi, and ECMr abundances within boundary spots. Error bars, 95% CI. Mann-Whitney U test (****P < 0.0001). UMAP, Uniform Manifold Approximation and Projection; NT, Normal Tissue; ST, Stomach Tumor; H&E, hematoxylin and eosin.

To investigate the histological spatial distribution of TME cell populations in their histologic context, we performed spatial transcriptomic analysis on two patients annotated as SFRP4-high GC (ST1 and ST2), a molecular subtype previously identified by our group ^23^ and one normal gastric section (NT1) obtained from the non-neoplastic region of a gastrectomy specimen (Method). Clinicopathologic characteristics of these patients are summarized in Table S1. NT1 displayed intact mucosal architecture with well-organized foveolar pits and glandular structures, a clearly delineated muscularis mucosae, and a loose connective tissue submucosa (Figure 1F and Figure S3A-S3C). In contrast, ST1 showed moderately differentiated tubular adenocarcinoma with infiltrative glandular structures surrounded by prominent desmoplastic stroma and scattered lymphoid aggregates, and the capture areas include the tumor invasive margins to characterize cellular organization at the interface of tumor progression (Figure 1G and Figure S3D-S3F). In addition, ST2 showed poorly differentiated adenocarcinoma with solid growth patterns and extensive desmoplastic stroma, and the capture area spanned from residual normal submucosa to the tumor mass, enabling characterization of the fibroblast composition gradient across the tumor-normal interface (Figure 1G, and Figure S3G-S3I). To spatially delineate the fibroblast subtypes, fibroblast subtype-informed deconvolution was performed using identified fibroblast subtypes except for the sample-specific MMP1+ fibroblasts. In NT1, SOX6+ fibroblasts were primarily localized within the mucosa layer (Figure S3C). Smooth muscle cells were specifically identified in the muscularis mucosae and pericytes were localized to vascular structures within the submucosa (Figure S3C). The submucosa was otherwise predominantly occupied by ECMh and ECMi subtypes (Figure 1F). In tumor samples, the stromal compartment was predominantly occupied by ECMr, in contrast to the ECMh-dominant stroma observed in NT1 (Figure 1G). In ST1, ECMr was enriched within the desmoplastic stroma surrounding infiltrative tumor, particularly at the invasion margin, whereas ECMi was preferentially distributed in stromal regions distal to the invasion margin and ECMh was sparsely detected (Figure 1G and Figure S3F). In ST2, the normal submucosa was dominated by ECMh, whereas the desmoplastic stroma adjacent to tumor nests was enriched for ECMr, with ECMi occupying an intermediate zone between the two regions (Figure 1G, Figure S3I). To determine which fibroblast subtype is spatially associated with malignant cells, we computed a gradient-based boundary score at the epithelial-stromal interface and quantified ECM fibroblast fractions within the boundary zone (Methods). Among the three ECM fibroblast subtypes, ECMr was the most enriched at the epithelial-stromal boundary, with significantly higher fraction compared with those of ECMh and ECMi (Figure 1H). These results indicate that ECMr fibroblasts preferentially localize to the tumor-stroma interface and represent the dominant ECM fibroblast subtype at the invasion front.

### Reprogramming of ECM fibroblasts in tumor microenvironment

The developmental relationships among ECM fibroblast subtypes were inferred from the trajectory analysis using partition-based graph abstraction (PAGA)^24^. The analysis revealed that the three ECM-related fibroblast subtypes formed a tightly connected subnetwork, suggesting potential developmental relationships among these fibroblast states (Figure 2A). Pseudotime ordering placed ECMh at the beginning of the inferred trajectory, followed by ECMi and then ECMr, positioning ECMi as an intermediate state and ECMr as a more differentiated state (Figure 2B and C, Table S4). Cellular differentiation potential was further evaluated using potency scores estimated by CytoTRACE2^25^, revealing that the ECMi cluster was notably enriched for predicted multipotent cells compared to other stromal populations (Figure 2D, Table S4). Quantitative comparison of CytoTRACE2 scores showed that ECMh and ECMi fibroblasts had similar median values, whereas ECMi fibroblasts exhibited a broader distribution with higher upper-range scores, suggesting greater developmental potential in a subset of ECMi fibroblasts (Figure 2E, left). Intriguingly, the predicted multipotent population was selectively expanded in tumors. Tumor-derived ECMh and ECMi fibroblasts displayed a markedly broader potency range, whereas their normal-derived counterparts remained in a low-potency state (Figure 2E, right). This high-potency state was also evident in the global UMAP distribution, which showed a continuum of fibroblast states between multipotent and more differentiated states (Figure 2F). Together, the concordance between pseudotime ordering and CytoTRACE2 potency scores supports a dynamic ECMh–ECMi–ECMr reprogramming trajectory characterized by the emergence of a predicted multipotent subset within ECMi, followed by progression toward a differentiated ECMr endpoint (Figure 2G).

**Figure 2.**
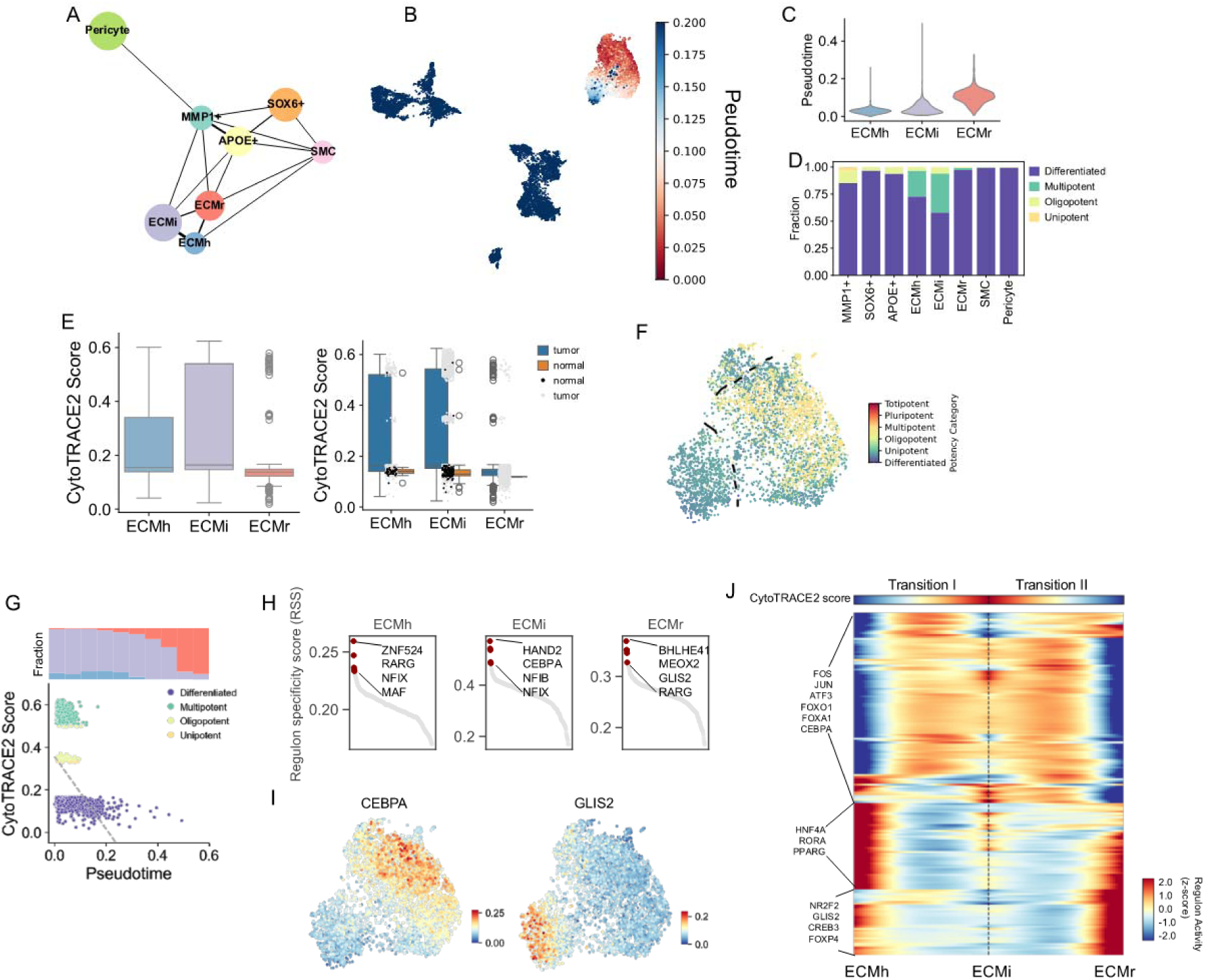
Reprogramming of ECM Fibroblasts in tumor microenvironment. (A) Partition-based graph abstraction (PAGA) of stromal populations. Edge thickness, PAGA connectivity. (B) UMAP colored by diffusion pseudotime with ECMh as the root. (C) Violin plot of pseudotime across ECM subsets. (D) CytoTRACE2 potency categories across stromal subsets. (E) CytoTRACE2 scores across ECM subsets. (Left) Pooled. (Right) Stratified by normal and tumor. Center line, median; box, IQR; whiskers, 1.5 × IQR. (F) Magnified ECM UMAP colored by CytoTRACE2 potency score. Dashed lines, manually delineated ECMh, ECMi, and ECMr territories. (G) (Top) Fractional composition of ECM subsets along pseudotime. (Bottom) CytoTRACE2 score versus pseudotime, colored by potency category. Dashed line, linear regression fit. (H) Ranked regulon specificity score (RSS) of SCENIC regulons across ECM subsets, with top 5 regulons per subset highlighted. (I) ECM UMAP colored by CEBPA and GLIS2 regulon activity (AUCell). (J) Heatmap of regulon activity along the CytoTRACE2-ordered branched trajectory, with ECMi treated as the shared branch point. Transition I, ECMh to ECMi; Transition II, ECMi to ECMr. Top track, CytoTRACE2 score. Values, z-scored GAM-smoothed AUCell activities. PAGA, partition-based graph abstraction; IQR, interquartile range; RSS, regulon specificity score; GAM, generalized additive model.

Our findings suggest that ECMi cells may acquire increased developmental potential within the tumor context and subsequently differentiate toward ECMr, a more differentiated tumor-associated fibroblast state. To identify the specific transcription factors (TFs) associated with the inferred reprogramming trajectory of ECM fibroblasts, we performed regulon analysis using SCENIC across all stromal populations (Figure S4A, Table S5). Regulon specificity score (RSS) analysis revealed that MEOX2 and GLIS2 ranked among the top-enriched regulons specific to ECMr, whereas CEBPA and HAND2 were preferentially associated with the ECMi state (Figure S4A and Figure 2H). Notably, these regulon assignments were specific to ECM fibroblast subtypes and were not shared with other stromal populations (Figure S4A). Visualization of regulon activities on the UMAP further showed that stress-responsive regulons, including ATF3, JUN, and FOS, were highly active in the predicted multipotent ECMi state, whereas GLIS2 regulon activity was concentrated in the ECMr subtype (Figure 2I and Figure S4B). To dissect the regulatory dynamics underlying ECM fibroblast reprogramming, we defined two distinct state transitions within the ECM fibroblast continuum. Transition I represented the transition from ECMh to the predicted multipotent ECMi state, whereas Transition II represented the subsequent transition from ECMi to the differentiated ECMr state. Branched expression analysis along CytoTRACE2 potency scores identified three regulon modules organized along the ECMh-ECMi-ECMr axis (Figure 2J, Table S6). One module exhibited progressively increased activity exclusively toward the ECMr state and included NR2F2, GLIS2, CREB3, and FOXP4 as candidate regulators of ECMr-state commitment (q-value < 2.0 × 10^−3^, Figure 2J). The other two modules captured regulons either peaking at the predicted multipotent ECMi intermediate or with reduced activity at ECMi relative to both differentiated endpoints (Figure 2J). Thus, ECM fibroblast reprogramming proceeds through a predicted multipotent ECMi intermediate followed by progression toward a differentiated ECMr remodeling state, with GLIS2 and its co-regulators emerging as candidate regulators of this transition.

### Spatial Characterization of Fibroblast Reprogramming in the Tumor Microenvironment

We delineated the spatial architecture of the tumor microenvironment into four distinct functional domains corresponding to epithelial, immune, stromal, and smooth muscle niches (ST1, Figure 3A and Figures S5A–S5C; ST2, Figure S6). We further investigated the distribution of fibroblast subtypes and observed spatial segregation between the ECMi and ECMr fibroblast fractions across the stromal domain (Figure 3B). Of note, we found that ECMi fibroblasts were preferentially localized to stromal regions outside the tumor margin, whereas ECMr fibroblasts were enriched at the tumor margin, forming an invasion-associated pattern adjacent to malignant niches (Figure 3B). Next, co-localization analysis revealed a negative spatial association between ECMi and ECMr (Moran’s I = −0.2778, p = 0.001, Figure 3C). To quantify the spatial proximity between spots or niches, we defined a tier distance metric as the minimum number of hexagonal grid steps on the Visium platform separating each spot from its nearest target boundary (Figure S5D, Methods). We then calculated the distance from each stromal spot to its nearest epithelial niche, which we termed ‘distance to cancer’ (Figure 3D). Across stromal spots, the ECMi fraction showed a significant positive correlation with distance to the epithelial niche (Pearson’s r = 0.591, p = 3.050 × 10^-67^, Figure 3E). In contrast, the ECMr fraction exhibited a significant negative correlation with this distance and increased progressively toward epithelial niches (r = −0.223, p = 2.383 × 10^-9^, Figure 3E). Accordingly, the relative contribution of ECMr within the combined ECMi–ECMr compartment increased in areas closer to the epithelial niche (Figure 3F).

**Figure 3.**
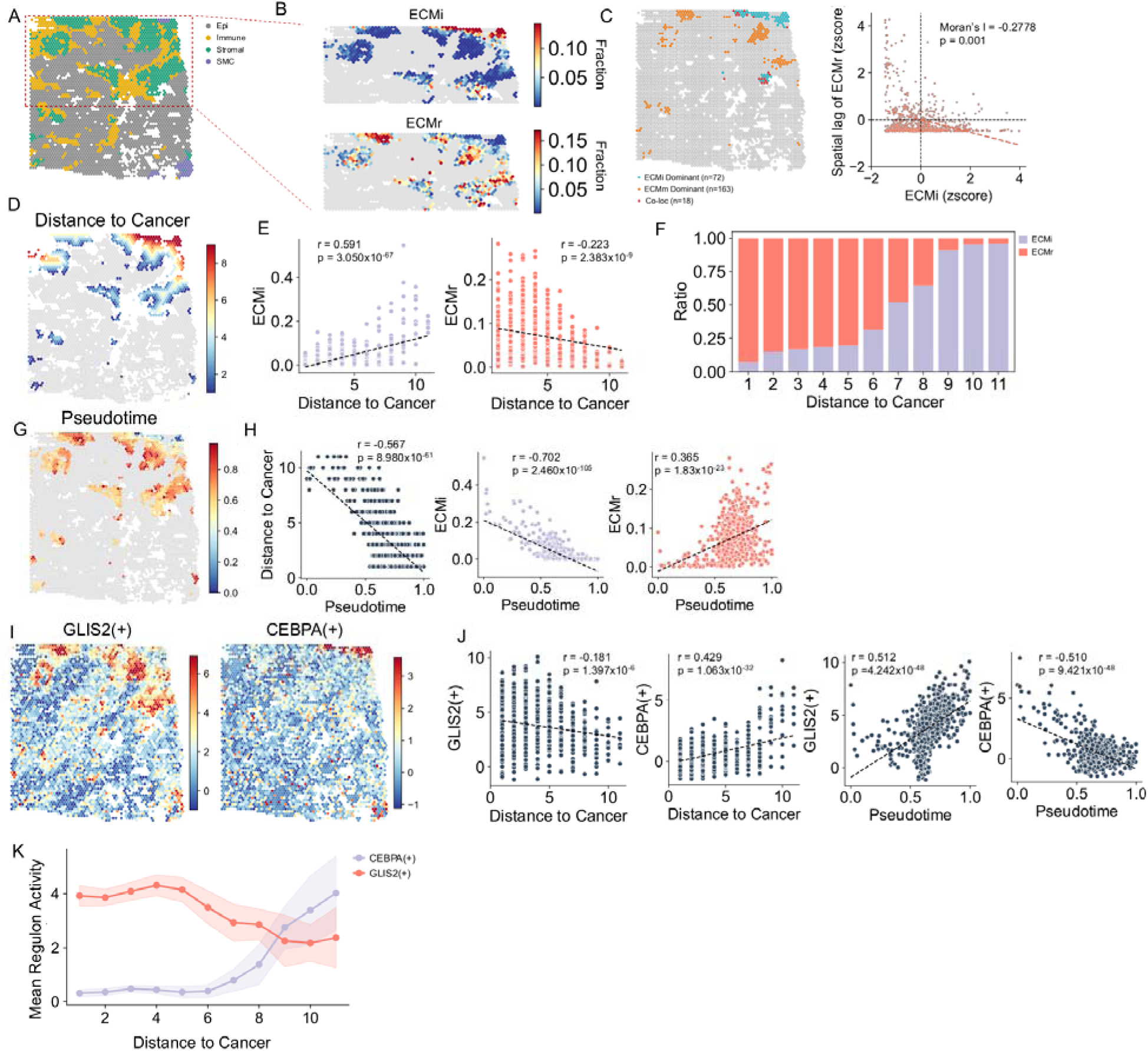
Fibroblast reprogramming in the tumor invasion margin. (A) Spatial niche annotation in ST1 by Local Indicators of Spatial Association (LISA) clustering. Red dashed box, region of interest shown in (B). (B) ECMi (top) and ECMr (bottom) fraction maps within the region of interest. (C) (Left) Spots classified by local bivariate Moran’s I into ECMi-dominant, ECMr-dominant, and co-localized categories (P < 0.05). (Right) Scatter of ECMi z-score against ECMr spatial lag, with global bivariate Moran’s I. (D) Distance to cancer map, defined as the minimum hexagonal grid steps from each stromal spot to the nearest epithelial spot. (E) ECMi (left) and ECMr (right) fractions against distance to cancer across stromal niche spots in ST1. Dashed lines, linear regression fit; Pearson r and P values are indicated. (F) Relative ECMi and ECMr composition across distance to cancer. (G) Diffusion pseudotime across stromal niche spots, with the cluster of highest ECMi fraction as the root. (H) Distance to cancer (left), ECMi (middle), and ECMr (right) fractions against pseudotime. (I) GLIS2(+) and CEBPA(+) regulon activity maps. Per-spot scores were inferred using the univariate linear model on the SCENIC-derived regulon network. (J) GLIS2(+) and CEBPA(+) activity against distance to cancer (left two) and pseudotime (right two). (K) Mean GLIS2(+) and CEBPA(+) regulon activity across distance to cancer bins. Shaded areas, 95% confidence intervals. LISA, Local Indicators of Spatial Association.

To spatially delineate the ECMi-to-ECMr transition, we performed spatial trajectory analysis within the stromal niche in ST1 (Figures 3 and S5), with corresponding analysis in ST2 (Figure S6). Based on the inferred connectivity among stromal states (Figures S5E–S5I), we calculated the diffusion pseudotime across the stromal niche (Figures S5E and S5G). Mapping the inferred trajectory onto the spatial coordinates revealed that spatially inferred pseudotime increased toward the tumor interface, showing a significant negative correlation with ‘distance to cancer’ (r = −0.567, p = 8.980 × 10^-61^; Figures 3G and 3H). This pseudotime gradient also demonstrated a significant negative correlation with the ECMi fraction and a significant positive correlation with the ECMr fraction (r = −0.702, p = 2.460 × 10^-105^ and r = 0.365, p = 1.83 × 10^-23^, respectively; Figure 3H), supporting the spatial alignment of the ECMi-to-ECMr transition relative to malignant cells. This spatial trajectory was further corroborated by the spatial activities of opposing transcription factor regulons. For instance, the GLIS2 regulon, associated with the differentiated ECMr endpoint, showed a significant negative correlation with the distance to epithelial spots and a positive correlation with spatial pseudotime (r = −0.181, p = 1.397 × 10^-6^ and r = 0.512, p = 4.242 × 10^-48^, respectively; Figures 3I and 3J). Conversely, the CEBPA regulon, associated with the ECMi state, displayed the opposite spatial gradient, showing a positive correlation with distance to cancer and a negative correlation with spatial pseudotime (r = 0.429, p = 1.063 × 10^-32^ and r = −0.510, p = 9.421 × 10^-48^, respectively; Figures 3I and 3J). When plotted along the distance from malignant spots, GLIS2 and CEBPA regulon activities exhibited a crossover pattern, delineating a spatial transition between ECMi-associated and ECMr-associated regulatory programs within the stromal niche (Figure 3K). This reciprocal spatial gradient of GLIS2 and CEBPA regulons further supports spatially organized ECMi-to-ECMr reprogramming along the invasive tumor front and its association with distinct transcription factor regulon activities.

### ECMr-associated GLIS2 regulon activity is linked to peritoneal recurrence

To evaluate the clinical significance of the ECMr-associated stromal remodeling program, we extended our analysis to two independent bulk-level microarray cohorts. Because ECMi and ECMr share core ECM marker expression, including *OGN*, *DCN*, and *LUM*, conventional bulk deconvolution approaches may not reliably distinguish these states. We therefore leveraged GLIS2 regulon activity as an ECMr-associated surrogate because it was preferentially enriched at the differentiated ECMr endpoint (Figure 2H) and exhibited a spatial gradient toward the tumor interface that paralleled ECMr enrichment (Figures 3I and 3J). We calculated GLIS2 regulon activity scores for two independent bulk-level microarray cohorts, the publicly available GSE84437 (n = 329) and ACRG cohort (GSE62254, n = 298) (Methods)^6,23,26^. First, when patients in GSE84437 and ACRG cohorts were stratified by median GLIS2 regulon activity, the GLIS2-high group exhibited significantly reduced overall survival in both cohorts (Figure S7A and S7B). To determine whether GLIS2 activity was associated with a specific route of postoperative metastatic spread, we compared GLIS2 regulon activity across metastasis types in GSE84437 (Figure S7C). Of note, GLIS2 activity was significantly elevated in patients with peritoneal metastasis compared to those with hematogenous (p = 6.316×10^-4^), lymph node (p = 1.262×10^-2^), or no recurrence (p = 3.081×10^-4^), whereas no significant difference was observed between peritoneal and combined peritoneal-hematogenous metastasis (p = 0.6256, Figure S7C). We next investigated the prognostic association between GLIS2 activity and peritoneal metastasis. Univariate Cox regression across metastasis types showed that GLIS2 activity was significantly associated with an increased risk of peritoneal metastasis (HR = 1.66, p = 5.280 × 10^-4^, Figure 4A), whereas no significant association was observed for hematogenous, lymph node, or other recurrence types (Figure 4A). Using the maximally selected rank statistic to optimize stratification for peritoneal metastasis-free survival (PMFS), Kaplan-Meier analysis demonstrated that GLIS2-high patients exhibited significantly reduced PMFS in both GSE84437 (p = 2.565 × 10^-4^, Figure 4B) and ACRG (p = 7.515 × 10^-13^, Figure 4C). Multivariate Co regression adjusting for T stage, N stage, sex, and age showed that the GLIS2-high group remained independently associated with reduced PMFS in both cohorts (HR = 2.12, p = 4.955×10^-3^ in GSE84437, Figure 4D; HR = 3.62, p = 2.9 × 10^-5^ in ACRG, Figure 4E). Importantly, GLIS2 regulon activity was measured in primary tumors obtained at surgery, before peritoneal metastasis became clinically apparent. Its association with subsequent PMFS therefore supports the interpretation that an ECMr-associated stromal remodeling program is already present in the primary tumor and marks tumors with a heightened propensity for peritoneal dissemination. To evaluate whether the association between GLIS2 activity and peritoneal metastasis extends beyond gastric cancer, we analyzed a colorectal cancer cohort (AMC-AJCCII-90, n = 86)^27,28^. GLIS2 regulon activity was significantly elevated in the CMS4 subtype compared to CMS1 and CMS2-3 subtypes (Figure S8A), consistent with the stromal-enriched nature of this molecular subtype. Kaplan-Meier analysis demonstrated that high GLIS2 activity was significantly associated with reduced peritoneal metastasis-free survival (p = 0.002, Figure S8B). These findings suggest that the link between GLIS2-associated stromal remodeling and peritoneal metastasis is not confined to gastric cancer but may represent a shared feature across gastrointestinal malignancies. Together, these findings identify GLIS2 regulon activity as an ECMr-associated primary-tumor stromal signature selectively linked to subsequent peritoneal metastasis in gastric cancer. To investigate potential communication between this clinically associated ECMr state and adjacent malignant cells at the invasion front, inferred cell-cell interactions between ECMr fibroblasts and malignant cells were next characterized.

**Figure 4.**
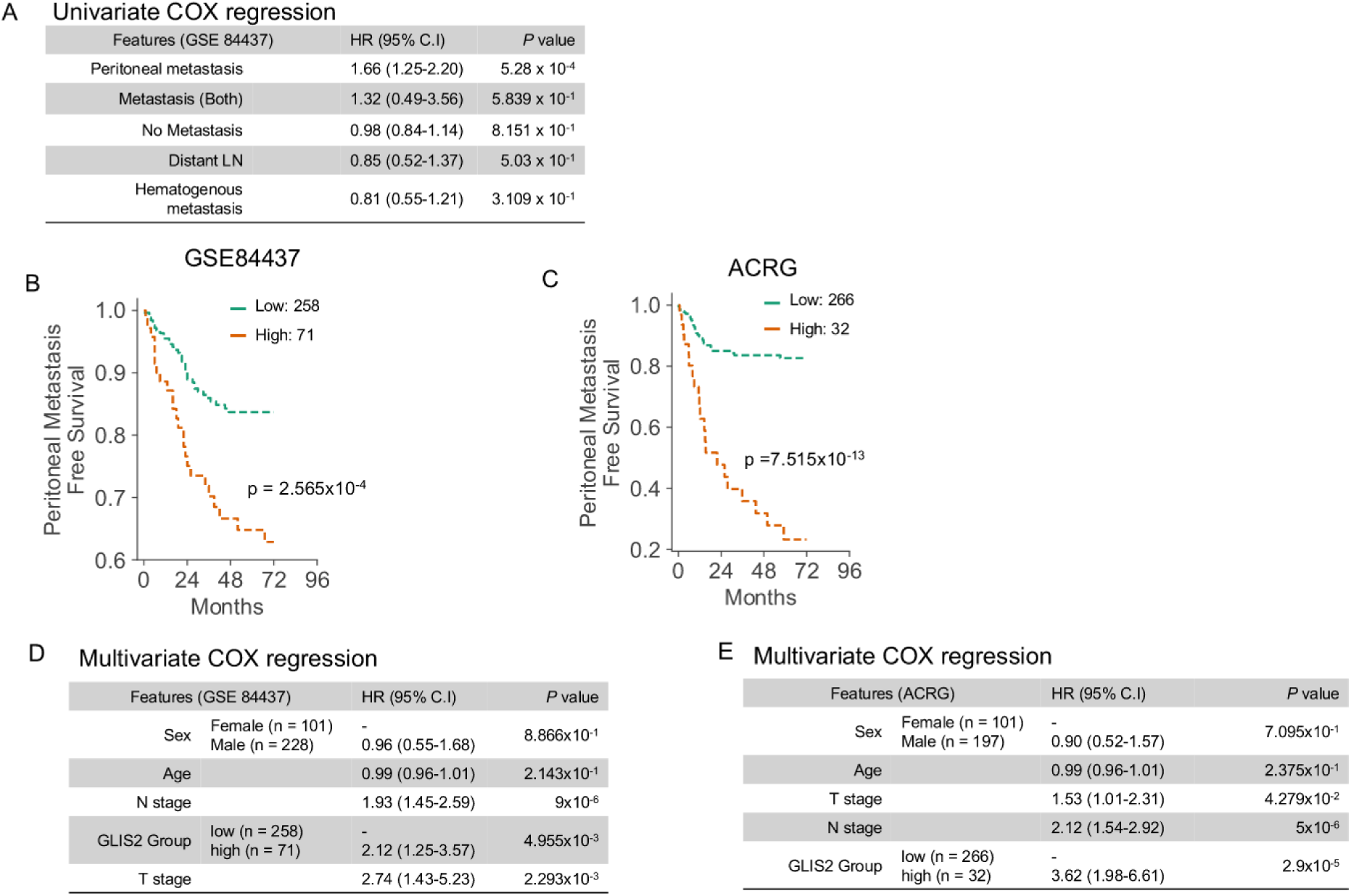
GLIS2 regulon activity is independently associated with peritoneal metastasis-free survival in gastric cancer. (A) Univariate Cox regression of GLIS2 regulon activity for recurrence types in the GSE84437 cohort. (B) Kaplan-Meier curves of peritoneal metastasis-free survival in the GSE84437 cohort, stratified into GLIS2(+) activity-high and -low groups by the maximally selected rank statistic. Log-rank test. (C) As in (B) for the ACRG cohort (GSE62254). (D) Multivariate Cox regression of peritoneal metastasis-free survival in the GSE84437 cohort, adjusted for sex, age, N stage, and T stage. (E) As in (D) for the ACRG cohort.

### Interaction inference nominates a reciprocal collagen–laminin axis at the tumor–stroma interface

To nominate candidate signaling axes underlying this interaction, cell-to-cell communication was inferred with CellChat. ECMr fibroblasts exhibited the second-highest outgoing signaling strength among all cell populations (Figure 5A), predominantly signaling through the COLLAGEN, LAMININ, and FN1 pathways (Figure 5B, Table S7). Focusing on interactions between ECMr fibroblasts and malignant cells, COLLAGEN and LAMININ emerged as the top reciprocal signaling axes, with COLLAGEN being the top ECMr-derived signal directed to malignant cells and LAMININ being the top malignant cell-derived signal directed to ECMr fibroblasts (Figure 5C). We next examined the dominant ligand-receptor pairs within the COLLAGEN and LAMININ axes. The analysis highlighted ligands produced by ECMr fibroblasts, including COL1A1 and COL1A2, and malignant cell-derived components of laminin-332 (LM332), comprising *LAMA3*, *LAMB3*, and *LAMC2* (Figure 5D). Notably, these LM332 components were predicted to interact with multiple receptors on ECMr fibroblasts, including the integrin complexes *ITGA1–ITGB1*, *ITGA9–ITGB1*, and *ITGAV–ITGB8*, as well as *CD44* (Figure 5D). Together, these inferred interactions nominate a reciprocal matrix-remodeling niche at the tumor–stroma interface. Given the dual role of LM332 in stable epithelial anchorage and migration during wound repair^29,30^, we proposed that fibroblast-derived collagen and malignant cell-derived LM332 constitute a reciprocal matrix circuit that repurposes an anchorage-associated interface into a dynamic, wound healing-like interface for epithelial migration and invasion.

**Figure 5.**
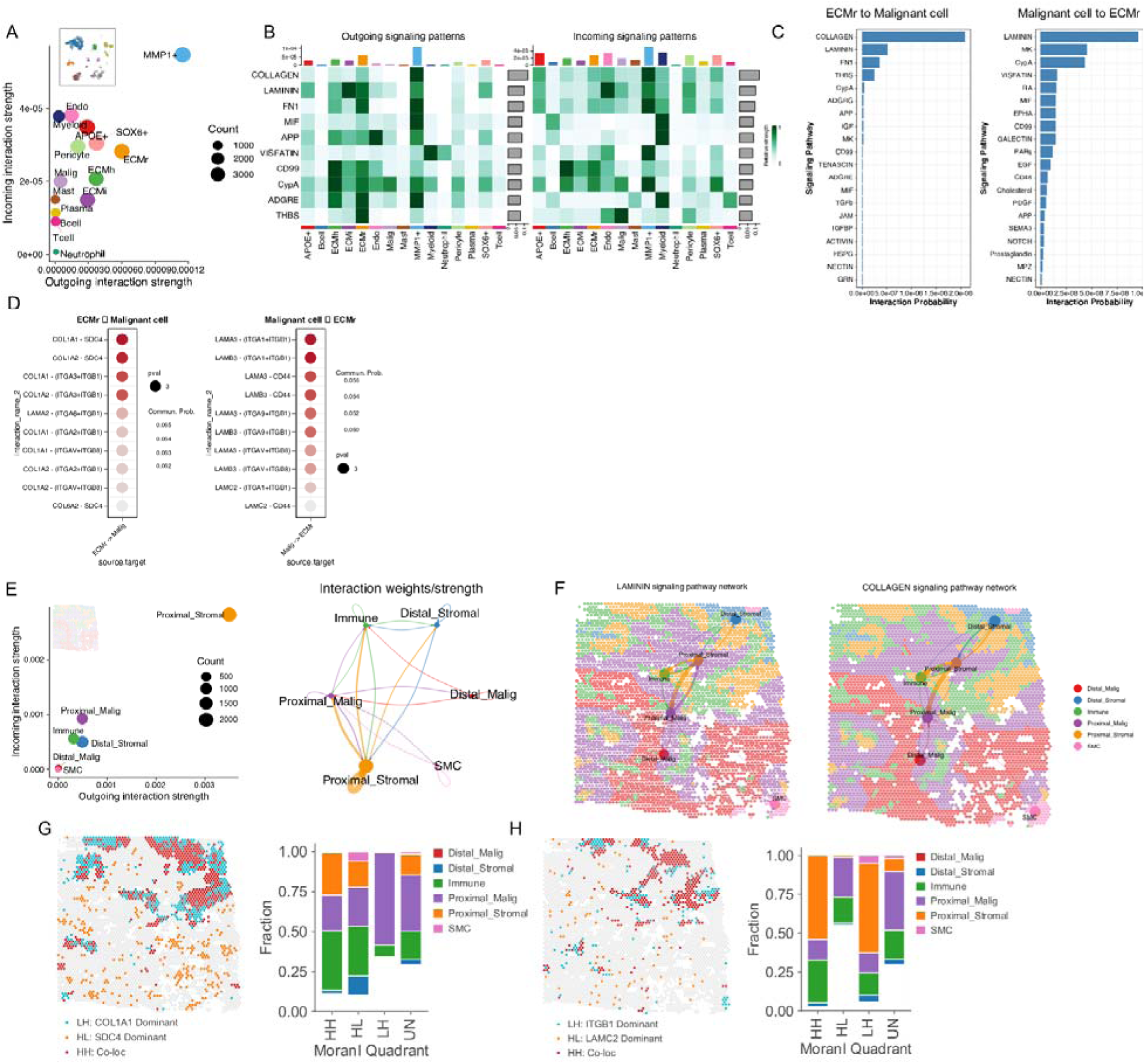
Reciprocal collagen-laminin signaling between ECMr fibroblasts and malignant cells at the tumor-stroma interface. (A) Outgoing and incoming interaction strength across cell populations inferred by CellChat in scRNAseq data. Dot size, number of inferred links. (B) Heatmaps of outgoing (left) and incoming (right) signaling patterns for the top 10 pathways. Top bars, relative signaling strength per pathway. (C) Top 20 signaling pathways ranked by interaction probability from ECMr to malignant cells (left) and malignant cells to ECMr (right). (D) Top 10 ligand-receptor pairs within COLLAGEN and LAMININ pathways for each direction. Color, communication probability; size, P value. (E) (Left) Interaction strength across six spatial niches. (Right) Interaction weight network in ST1. (F) LAMININ (left) and COLLAGEN (right) signaling pathway networks in ST1. Spot color, niche; node size, spots per niche; edge thickness, interaction strength. (G) (Left) Local bivariate Moran’s I map of COL1A1 and SDC4 in ST1, classified into HH (co-localized), HL (SDC4-dominant), and LH (COL1A1-dominant). (Right) Niche composition per quadrant. UN, non-significant. (H) As in (G) for LAMC2 and ITGB1.

We next examined the spatial organization of these inferred interactions in two spatial transcriptomic datasets (ST1, Figures 5 and S9; ST2, Figure S10). The spatial relationship between epithelial and stromal niches was defined based on the distance metric described above, with spots within five hexagonal steps of the epithelial–stromal boundary categorized as proximal (Figures S9A and S10A). Spatial niche-level cell-to-cell interaction analysis across six spatially defined niches identified the proximal stromal–malignant interface as exhibiting the highest inferred interaction frequency and overall signaling weight (Figures 5E, S9B, and S10B). Consistent with the single-cell results, the COLLAGEN, FN1, and LAMININ pathways showed the highest inferred information flow between the proximal stromal and malignant niches (Figure S9C). Spatial mapping of interaction strength showed that the inferred COLLAGEN and LAMININ signals were preferentially concentrated at the proximal stromal–malignant boundary (Figures 5F, S9D, and S9E). Co-localization analysis of key ligand-receptor pairs further demonstrated that COL1A1–SDC4 and LAMC2–ITGB1 were co-enriched within the proximal stromal niche (Figures 5G, 5H, S10F, and S10G), supporting the spatial convergence of key ligand-receptor components of the reciprocal collagen–laminin axis at the tumor–stroma interface. Collectively, these findings spatially corroborate the reciprocal collagen–laminin matrix circuit nominated by the single-cell analysis. We next evaluated whether ECMr fibroblasts induce the LM332- and hemidesmosome-associated wound healing-like program predicted by this model in malignant cells.

### *In vitro* validation of ECMr-mediated transcriptional reprogramming in cancer cells

We performed in vitro validation experiments to evaluate the functional impact of ECMr fibroblasts on cancer cells. We immortalized and tdTomato-labeled patient-derived fibroblasts (pdFib) and established four monoclonal fibroblast lines (Methods, Figure 6A, and Figure S11A). Microscopic examination revealed that the parental immortalized pdFib pool and all four monoclonal cell lines exhibited a myofibroblast-like morphology (Figure S11B). To validate their ECM subtype identity, signature scores derived from scRNA-seq markers confirmed that the parental pool and all four monoclonal lines corresponded to the ECMr state (Figure 6B, Methods). The four clones additionally exhibited higher GLIS2 regulon activity than the parental pool (Figure 6C). We then conducted a 14-day co-culture experiment by culturing each monoclonal ECMr fibroblast line with GFP-labeled SNU668 gastric cancer cells (Figure 6A and Figure S11C).

**Figure 6.**
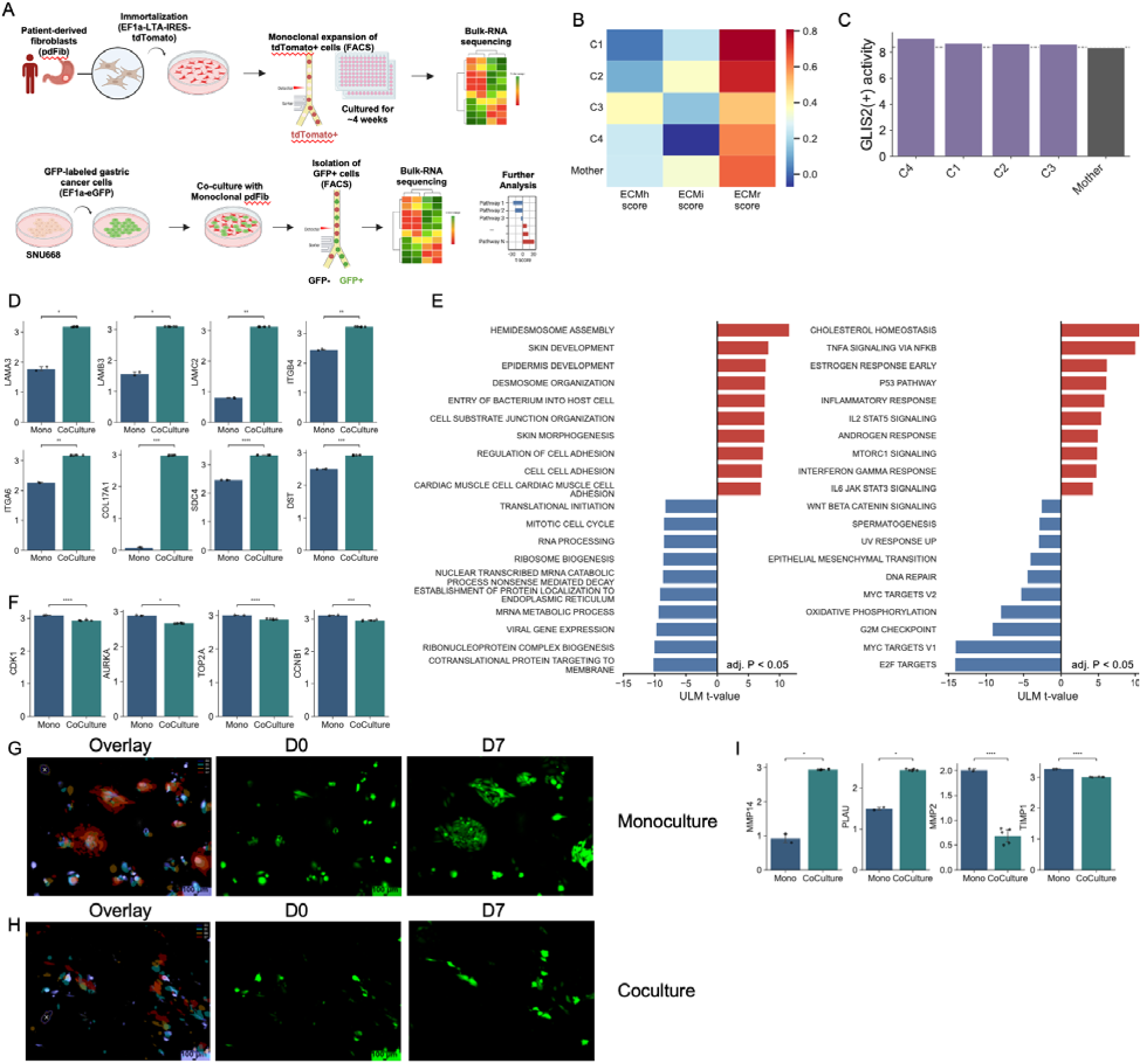
In vitro validation of ECMr-induced reprogramming in malignant cells. (A) Schematic of the experimental workflow. (B) Heatmap of ECMh, ECMi, and ECMr signature scores in four monoclonal pdFib lines (C1 to C4) and the parental pool (Mother). (C) GLIS2(+) regulon activity in monoclonal pdFib lines and the parental pool. Dashed line, parental pool activity. (D) Bar plots of LM332- and hemidesmosome-associated gene expression in 14-day time-matched monoculture controls (Mono, n = 2) and 14-day ECMr co-cultured SNU668 cells (n = 5; C2, n = 1; C3, n = 2; C4, n = 2). Y axis, log2 TPM. Error bars, 95% confidence intervals. Welch’s t-test (*P < 0.05, **P < 0.01, ***P < 0.001, ****P < 0.0001). (E) Top 10 up- and down-regulated GO Biological Process (left) and Hallmark (right) pathways in co-cultured versus monoculture SNU668 cells, ranked by ULM t-value applied to PyDESeq2 Wald statistics. (F) As in (D) for cell cycle genes. (G) Longitudinal imaging of SNU668 cells in monoculture by fluorescence inverted microscopy. (Left) Overlay of GFP signal at D0, D1, D4, and D7, with each time point assigned a distinct color. (Middle and right) GFP signal at D0 and D7. Scale bar, 100 µm. (H) As in (G) for SNU668 cells co-cultured with ECMr fibroblasts. (I) As in (D) for protease genes.

Following 14 days of co-culture, GFP-positive cancer cells were isolated by FACS and subjected to RNA sequencing to identify transcriptional changes induced by ECMr fibroblast co-culture. Post-sort purity assessment by re-gating the sorted samples revealed 3.5% residual fibroblast contamination in the clone 1 co-culture sample, accompanied by an elevated EMT score relative to the other co-culture samples (Figures S11D and S11E). Accordingly, the clone 1 co-culture sample was excluded on the basis of residual fibroblast contamination, and the primary transcriptomic analysis compared 14-day time-matched monoculture controls with cancer cells co-cultured for 14 days with the remaining three clones, whose sorted cancer cell populations showed high purity (Figure S11D). Compared with the time-matched monoculture controls, cancer cells co-cultured with ECMr fibroblasts showed significant upregulation of *LAMC2*, *LAMA3*, and *LAMB3*, providing experimental support for the malignant-cell origin of the LM332 signal nominated by CellChat (Figure 6D, Table S8). Pathway activity analysis identified hemidesmosome assembly as the top-ranked positively enriched gene set (Figure 6E, left). Consistent with this enrichment, additional adhesion-associated genes were significantly upregulated, including the α6β4 integrin subunits *ITGA6* and *ITGB4*, the transmembrane hemidesmosomal protein *COL17A1*, the cytoskeletal linker *DST*, and the ECM co-receptor *SDC4* (Figure 6D). Together, these changes define a coordinated LM332–hemidesmosome-associated transcriptional program encompassing matrix ligand production, adhesion receptors, and hemidesmosomal structural components.

Examining the suppressed gene programs, Hallmark pathway analysis revealed that MYC targets, E2F targets, and G2M checkpoint were the most strongly downregulated pathways (Figure 6E, right), with concordant reductions in the inferred activities of MAPK, PI3K, and EGFR signaling (Figure S12A). Consistent with these pathway-level changes, individual cell cycle genes including CDK1, TOP2A, AURKA, and CCNB1 were each significantly downregulated in cancer cells co-cultured with ECMr fibroblasts relative to the time-matched monoculture controls (Figure 6F), and broader examination showed that the majority of cell cycle-related genes were more highly expressed in monoculture controls than in co-cultured cancer cells (Figure S12B). To assess proliferative behavior under co-culture conditions, GFP-labeled SNU668 cancer cells were seeded on top of mitomycin C-treated ECMr monoclonal fibroblasts and tracked longitudinally over 7 days (Figures 6G, 6H, S12D, and S12E). Compared with monoculture, in which cancer cells progressively proliferated and formed expanding clusters at their initial positions, co-cultured cancer cells showed minimal cluster expansion throughout the imaging period. Notably, co-cultured cells instead exhibited progressive morphological elongation and positional displacement, in contrast to the spatially stable, compact clusters observed in monoculture, with the same pattern reproduced in an independent experiment (Figures 6G, 6H, and S12D–S12G). These observations led us to consider whether the malignant cell-derived LM332 signal nominated by single-cell analysis and the LM332–hemidesmosome-associated transcriptional program induced in vitro might represent a wound healing-like program rather than static cell anchoring. To probe this possibility, we examined proteases relevant to wound re-epithelialization.

*MMP14* (MT1-MMP) *and PLAU* (uPA), both implicated in pericellular matrix remodeling and keratinocyte migration during wound re-epithelialization, were significantly upregulated, whereas *MMP2* and *TIMP1* were significantly downregulated (Figure 6I). In addition, the Hallmark epithelial–mesenchymal transition pathway was significantly downregulated in co-cultured cancer cells (Figure 6E, right, and Figure S11E). Transcription factor activity analysis further supported this observation, showing increased inferred activities of the epithelial identity regulators CDX2 and GRHL2, together with reduced activities of the EMT regulators ZEB1 and SNAI2 and the proliferative regulator MYC (Figure S12C). Together, these results indicate that ECMr co-culture induces a wound healing-like epithelial reprogramming state characterized by reduced proliferation and transcriptionally distinct from classical EMT.

### Spatial transcriptomics reveals a niche-associated Go-or-Grow architecture at the tumor–stroma interface

We next investigated whether these co-culture-induced programs were spatially recapitulated in primary GC tissues. Given that ECMr co-culture induced coordinated upregulation of all three LM332 subunit genes in cancer cells (Figure 6D), we first examined the spatial distribution of *LAMC2*, one of the most significantly upregulated genes following co-culture. In ST1, *LAMC2* expression peaked at the stromal–epithelial boundary and progressively declined with increasing distance into the epithelial niche (Figures 7A and 7B). The expression of *LAMA3* and *LAMB3* exhibited the same spatial gradient, peaking at the invasion front and declining toward the distal tumor bulk (Figures S13A and S13B), supporting the coordinated spatial expression of all three LM332 subunit genes at the tumor–stroma interface. This spatial pattern indicates that the LM332-associated transcriptional program is concentrated at the tumor invasion front, consistent with the reciprocal signaling architecture nominated by CellChat analysis (Figure 5).

**Figure 7.**
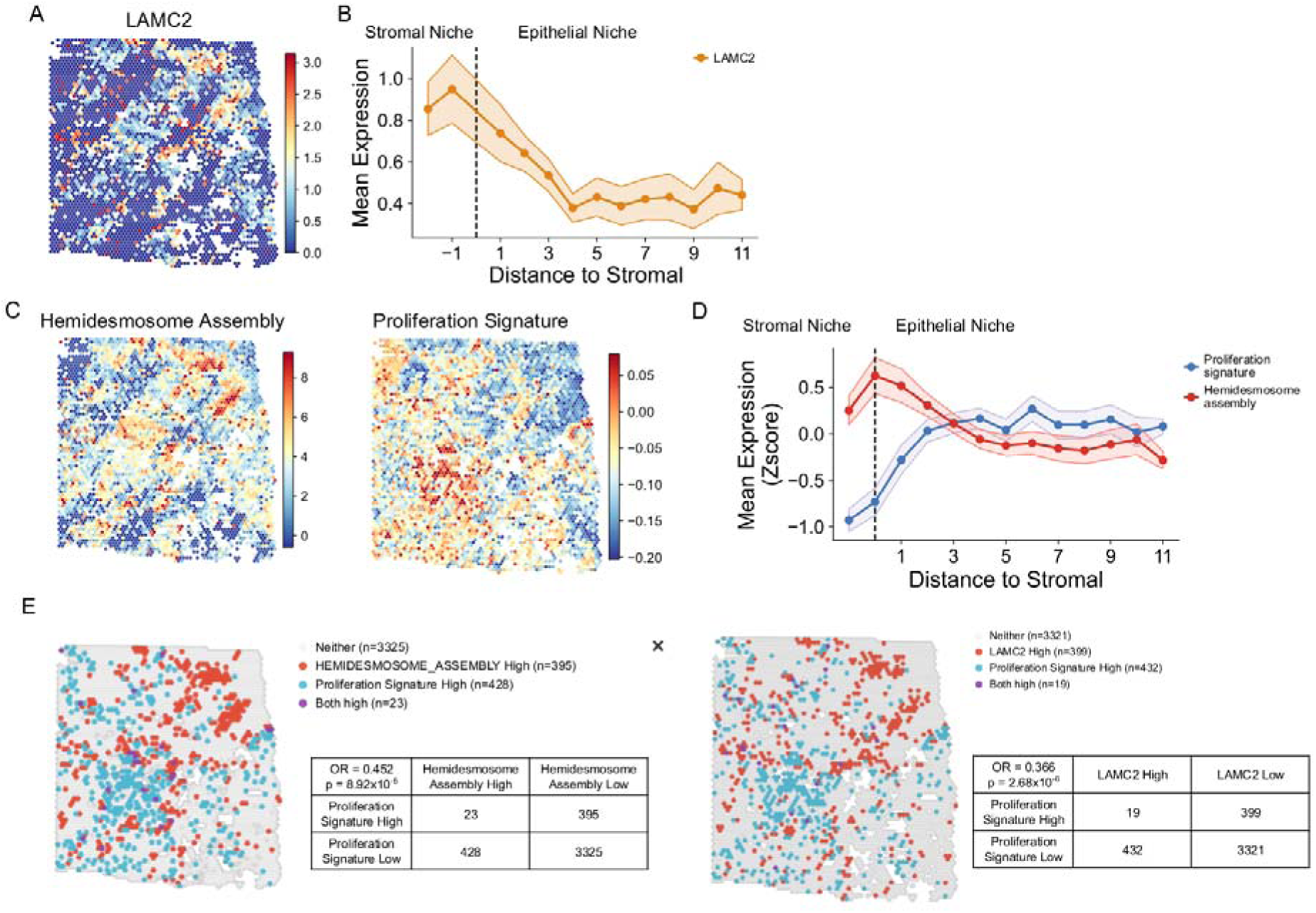
Spatial transcriptomics reveals a niche-associated Go-or-Grow architecture at the tumor–stroma interface. (A) Spatial map of LAMC2 expression in ST1. Color, log-normalized expression. (B) Mean LAMC2 expression across distance to stromal from the tumor–stroma interface; negative and positive values denote the stromal and epithelial sides, respectively in ST1. Dashed line, stromal-epithelial boundary. Shaded area, 95% confidence interval. (C) Spatial maps of hemidesmosome assembly (left) and proliferation signature (right) scores in ST1. (D) Mean proliferation signature and hemidesmosome assembly scores (z-scored) across distance to stromal in ST1. Dashed line, stromal-epithelial boundary. Shaded areas, 95% confidence intervals. (E) (Left) Spatial map of ST1 with epithelial niche spots classified by hemidesmosome assembly (90th percentile threshold) and proliferation signature (zero threshold) scores. Associated 2 × 2 contingency table with Fisher’s exact test. (Right) As in (left) for LAMC2 expression and proliferation signature.

We next scored the proliferation signature and the hemidesmosome assembly gene set (Methods) and plotted their activities along the tumor–stroma axis (Figure 7C). The two programs exhibited reciprocal spatial gradients, with the hemidesmosome assembly score peaking at the tumor invasion front and proliferation increasing toward the distal tumor bulk (Figure 7D). To quantify this spatial segregation, we performed co-localization analysis by categorizing malignant-niche spots based on their proliferation and hemidesmosome assembly activities. The two programs exhibited a significant negative spatial association, with relatively few spots classified as high for both programs (OR = 0.452, p = 8.92 × 10^-5^; Figure 7E). This negative association was recapitulated at the single-gene level, with LAMC2-high and proliferation-high spots showing a comparable spatial exclusion pattern (OR = 0.366, p = 2.68 × 10^-6^; Figure 7E). The agreement between pathway- and single-gene-level analyses, together with robustness across a range of classification thresholds, supports a spatially resolved inverse relationship between the hemidesmosome-associated transcriptional program and proliferation at the tumor–stroma interface (Figure S13C).

Given the matrix-remodeling protease pattern associated with the wound healing-like state observed in vitro, characterized by decreased *MMP2* and increased *MMP14* and *PLAU* (Figure 6I), we next examined whether this pattern was spatially recapitulated. In ST1, the hemidesmosome assembly score was negatively associated with MMP2 (OR = 0.645, p = 2.55 × 10^-2^; Figure S13D). In contrast, the hemidesmosome assembly score significantly co-localized with *PLAU* (OR = 1.968, p = 6.55 × 10^-6^), whereas its association with *MMP14* showed a concordant but non-significant trend (OR = 1.315, p = 0.086; Figure S13E). Malignant-niche spots proximal to ECMr-enriched stroma exhibited elevated LM332-subunit expression and hemidesmosome assembly scores together with the associated protease pattern, whereas proliferative activity was enriched in more distal tumor regions. These spatial patterns were recapitulated in the independent ST2 specimen (Figure S14). Together, the concordance between the co-culture and two spatial datasets supports an ECMr-associated Go-or-Grow architecture in which the hemidesmosome-associated wound healing-like program is enriched at the tumor–stroma interface, whereas proliferation predominates in more distal tumor regions.

### Recapitulation of ECM fibroblast spatial architecture and ECMr–tumor coupling in peritoneal metastasis

Based on the clinical association between GLIS2 activity and peritoneal metastasis, we next examined whether the stromal remodeling and tumor–stroma interaction programs identified in primary tumors were recapitulated at the peritoneal metastatic site. We analyzed spatial transcriptomics data from a peritoneal metastasis sample (PT1)^31^, in which the captured tissue spanned from the peritoneal surface through tumor-infiltrated stroma to the underlying fat tissue (Figure 8A). Spatial deconvolution revealed that epithelial and ECMr fractions were co-distributed across tumor-involved regions, whereas ECMi occupied spatially distinct regions (Figures 8B and S15). Niche analysis delineated four spatial domains corresponding to epithelial, immune, stromal, and smooth-muscle compartments, as observed in the primary tumors (Figure 8C). Concordantly, bivariate Moran’s I analysis demonstrated a strong positive spatial correlation between ECMr and epithelial fractions (global Moran’s I = 0.511, p = 0.001; Figure 8D), in contrast to the primary tumors, where ECMr enrichment was concentrated at the tumor–stroma boundary.

**Figure 8.**
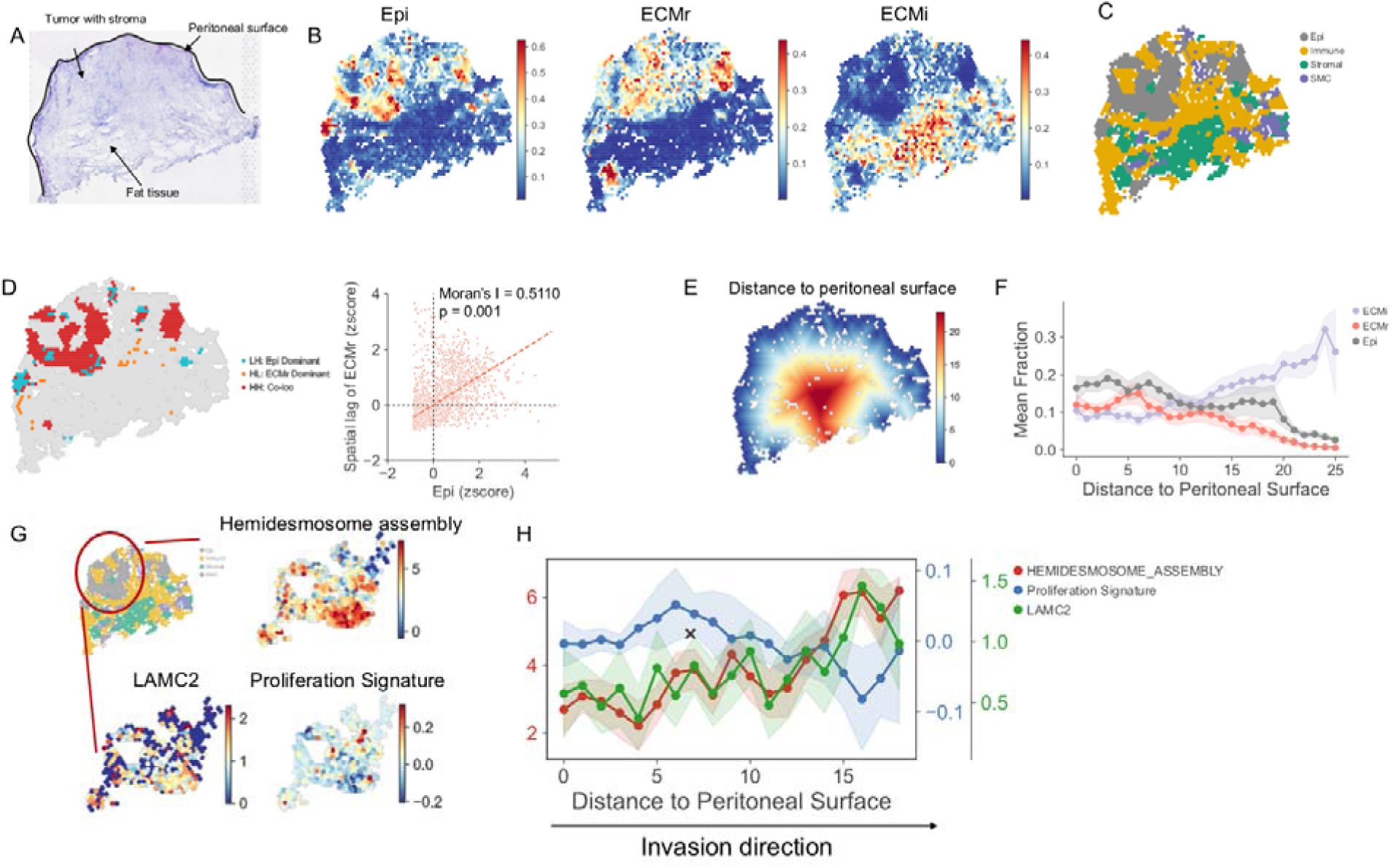
Stromal architecture and the wound healing-like program are recapitulated in peritoneal metastasis. (A) H&E image of the peritoneal metastasis tissue (PT1) with manual annotation of the peritoneal surface. (B) Spatial maps of Epi, ECMr, and ECMi fractions in PT1. (C) Spatial niche annotation in PT1 by LISA clustering. (D) (Left) Local bivariate Moran’s I spatial map of Epi and ECMr fractions in PT1. (Right) Scatter plot of Epi z-score against the spatial lag of ECMr z-score, with global bivariate Moran’s I. (E) Spatial map of distance to peritoneal surface in PT1. (F) Mean fractions of Epi, ECMr, and ECMi across distance to peritoneal surface in PT1. Shaded areas, 95% confidence intervals. (G) Spatial maps of LAMC2 expression, hemidesmosome assembly score, and proliferation signature score within an epithelial niche region of interest in PT1. Red circle in the inset indicates the region of interest. (H) Mean hemidesmosome assembly score, proliferation signature score, and LAMC2 expression across distance to peritoneal surface within the region of interest in (G). Arrow indicates the histologically inferred direction of tumor extension. Shaded areas, 95% confidence intervals.

To examine whether the proximal ECMr–distal ECMi organization observed in primary tumors was preserved in the metastatic tissue, we used the histologically defined peritoneal surface as the origin of a tissue-depth axis corresponding to the inferred direction of tumor extension into deeper tissue layers and calculated the distance of each spot from this surface (Figures 8E and S16A). We then compared ECM fibroblast fractions along this axis (Figure 8F). The ECMr fraction was highest near the peritoneal surface and gradually declined toward deeper layers, whereas the ECMi fraction exhibited the opposite trend, being lowest near the surface and progressively increasing with tissue depth (Figure 8F). The epithelial fraction was likewise enriched near the peritoneal surface and declined at greater tissue depths, broadly paralleling the ECMr distribution (Figure 8F). This spatial organization recapitulates the fundamental relationship observed in primary tumors, where ECMr fibroblasts were enriched proximal to malignant cells and ECMi predominated in more distal stromal regions.

We next examined whether the hemidesmosome-associated gene program and Go-or-Grow spatial architecture were also present in the metastatic tissue. To this end, we defined a region of interest within the epithelial niche of PT1 and quantified the hemidesmosome assembly score, proliferation signature, and *LAMC2* expression, which had shown coordinated spatial organization in the primary tumors. Within this epithelial niche, the hemidesmosome assembly and proliferation scores displayed spatially distinct distributions (Figures 8G and S16B), consistent with the inverse relationship observed at the primary tumor invasion front (Figure 7). LAMC2 expression exhibited a spatial pattern concordant with the hemidesmosome assembly score, with both enriched toward the histologically inferred deeper invasion front (Figure 8G). When plotted along the distance-from-the-peritoneal-surface axis, the hemidesmosome assembly score and *LAMC2* expression were lowest near the peritoneal surface and progressively increased toward the deeper invasion front, whereas the proliferation signature showed the opposite trend, with the highest activity near the surface and declining with tissue depth (Figures 8H and S16C). The three spatial profiles exhibited a crossover at the mid-depth region, recapitulating the Go-or-Grow spatial architecture observed in primary tumors. Together, these findings show that the ECMi–ECMr spatial segregation and the inverse spatial organization of the hemidesmosome-associated and proliferative programs observed in primary gastric tumors were recapitulated in PT1, together with widespread ECMr–epithelial spatial coupling across tumor-involved regions.

## Discussion

In this study, we delineated a spatially organized continuum of extracellular matrix (ECM)-producing fibroblast states in gastric cancer by integrating single-cell transcriptomics, spatial transcriptomics, patient-derived fibroblast co-culture, and clinical cohort analyses. We identified three transcriptionally distinct fibroblast states—ECMh, ECMi, and ECMr—ordered along an inferred continuum, with the tissue-remodeling ECMr state and GLIS2 regulon activity preferentially enriched near the tumor–stroma interface. Across two independent gastric cancer cohorts, ECMr-associated GLIS2 regulon activity was independently associated with shorter peritoneal metastasis-free survival, linking this ECMr-associated stromal program to subsequent peritoneal dissemination. Cell–cell interaction analysis nominated reciprocal collagen–laminin matrix signaling between ECMr fibroblasts and malignant cells, and co-culture with patient-derived ECMr fibroblasts elicited a non-EMT epithelial transcriptional program characterized by coordinated LM332-subunit expression, a hemidesmosome-associated gene program, a matrix-remodeling protease profile, and reduced proliferative activity. Longitudinal imaging further showed progressive elongation and positional displacement of cancer cells over the ECMr fibroblast layer. This epithelial program was enriched at the tumor–stroma interface and inversely organized with proliferation across the spatial datasets. ECMi–ECMr spatial segregation, widespread ECMr–epithelial spatial coupling, and related epithelial organization were also observed in a peritoneal metastasis. Together, these findings define an ECMr-centered tumor–stroma ecosystem in primary gastric cancer that is associated with subsequent peritoneal dissemination.

The ECM fibroblast continuum identified in this study supports the view that fibroblast heterogeneity in gastric cancer reflects dynamic state transitions rather than fixed, unrelated subtypes. ECMh fibroblasts were preferentially represented in non-neoplastic tissue, whereas ECMi and ECMr states were expanded in tumors. ECMi fibroblasts exhibited a broader CytoTRACE2 score distribution extending into higher predicted potency values and occupied an intermediate position along the inferred trajectory, whereas ECMr fibroblasts showed a more differentiated transcriptional profile and localized preferentially near the tumor–stroma interface. These observations support a two-stage reprogramming model in which tumor-associated cues shift ECMh fibroblasts toward a transcriptionally plastic ECMi intermediate, followed by transition to the tissue-remodeling ECMr state in proximity to malignant cells. This interpretation remains inferential because lineage relationships were reconstructed computationally rather than demonstrated by direct lineage tracing. Nevertheless, the concordance among pseudotime, predicted potency, regulon activity, and spatial gradients, together with the recovery of all three states in two independent scRNA-seq cohorts, supports the proposed ECMh–ECMi–ECMr reprogramming trajectory in the gastric tumor microenvironment.

Regulon analysis further distinguished the ECM fibroblast states. CEBPA-associated activity was enriched in the ECMi intermediate, whereas GLIS2 regulon activity increased toward the ECMr-associated end of the trajectory and was spatially enriched near the tumor–stroma interface. These data position GLIS2 regulon activity as a reproducible transcriptional feature of the ECMr state, but they do not establish GLIS2 itself as a functional driver of fibroblast reprogramming. Accordingly, GLIS2 regulon activity serves here as an analytical surrogate for estimating ECMr-associated stromal activity in bulk transcriptomic cohorts. Direct perturbation experiments will be required to determine whether GLIS2 contributes causally to the acquisition or maintenance of the ECMr phenotype.

A major observation of this study is that ECMr fibroblast co-culture induced a hemidesmosome-associated transcriptional program in cancer cells, with hemidesmosome assembly emerging as the top-ranked positively enriched gene set. Co-cultured cancer cells upregulated all three laminin-332 (LM332) subunit genes, LAMA3, LAMB3, and LAMC2, together with the α6β4 integrin subunits ITGA6 and ITGB4 and the transmembrane hemidesmosomal component COL17A1. Hemidesmosomes have traditionally been viewed as stable epithelial anchoring structures whose dissolution facilitates invasion^32,33^. However, transient or nascent hemidesmosome-like adhesions can also support epithelial migration during wound re-epithelialization^29,30,34^. In this setting, migrating keratinocytes deposit laminin-332 and dynamically assemble adhesion complexes that provide traction while preserving epithelial identity. Together, this transcriptional response more closely resembles a wound healing-like epithelial migratory state that preserves epithelial identity than canonical EMT.

Importantly, the wound-repair parallel extends beyond the epithelial response to the fibroblast compartment itself. During physiological wound repair, resident fibroblasts depart from homeostasis and acquire spatially and temporally organized inflammatory and matrix-remodeling states^19,20^. The spatially organized ECMh–ECMi–ECMr reprogramming trajectory toward the tumor invasion margin recapitulates this stromal logic, while malignant cells at the interface engage an LM332–hemidesmosome-associated program resembling that used by wound-edge epithelia^29,30,34^. Previous study performed by Hu et al. identified coordinated multicellular gene programs shared between healing wounds and tumors^35^. Our findings extend this concept by resolving a specific stromal–epithelial configuration at the gastric cancer interface, in which ECMr-associated matrix remodeling converges with an LM332–hemidesmosome-associated epithelial response. This convergence supports a model in which gastric cancer co-opts a coordinated stromal–epithelial wound-repair circuit, creating a persistent wound-like interface that supports epithelial movement and continued matrix remodeling.

This wound-repair interpretation is further supported by the accompanying protease profile. ECMr-exposed cancer cells upregulated MMP14 and PLAU, which participate in pericellular matrix remodeling during epithelial migration and wound re-epithelialization, while MMP2 and TIMP1 were reduced^36–38^. Because both MMP14 and MMP2 can contribute to LM332 processing and matrix remodeling, this expression pattern does not identify a single LM332 cleavage mechanism. Rather, it indicates a reconfigured pericellular protease expression program accompanying the LM332–hemidesmosome-associated response. Taken together, the induction of LM332 and hemidesmosome-associated components alongside this matrix-remodeling protease shift suggests that cancer cells at the fibroblast-rich interface may adopt an epithelial migration program adapted to a remodeled extracellular matrix. These findings do not demonstrate that mature hemidesmosome structures form or are required for migration, but they provide a molecular framework for subsequent functional investigation.

The co-culture and spatial analyses also revealed an inverse relationship between the hemidesmosome-associated program and proliferation. Cancer cells exposed to ECMr fibroblasts showed suppression of MYC targets, E2F targets, G2M checkpoint activity, and multiple cell-cycle genes. Longitudinal imaging showed progressive elongation and positional displacement of cancer cells over the ECMr fibroblast layer, with minimal apparent expansion of tumor-cell clusters. This behavior is consistent with a fibroblast-supported mode of epithelial movement on a living stromal substrate rather than proliferation-driven cluster expansion. In the spatial datasets, LM332-subunit expression and the hemidesmosome-associated program peaked at the tumor–stroma interface, whereas proliferation increased toward the tumor interior. Although these spatial data do not directly resolve temporal switching between cellular states, the reciprocal organization is consistent with a Go-or-Grow-like spatial architecture in which a matrix-engaged, less proliferative epithelial program predominates within an ECMr-associated niche.

Importantly, this tumor-cell program was observed without activation of canonical EMT. ECMr co-culture suppressed EMT-associated pathways and was associated with reduced inferred activity of EMT-related regulators, whereas epithelial identity regulators such as CDX2 and GRHL2 showed increased inferred activity. These results are relevant to the interpretation of stromal-rich molecular subtypes, including the ACRG MSS/EMT subtype in gastric cancer and CMS4 in colorectal cancer. Bulk EMT signatures can incorporate substantial stromal expression and thereby confound estimates of tumor cell-intrinsic mesenchymal transition^6–8^. In the present study, residual fibroblast contamination in one sorted co-culture sample was accompanied by an elevated EMT score, illustrating the sensitivity of bulk or incompletely resolved data to stromal admixture. Our findings therefore support a model in which stromal-rich gastric cancers can exhibit invasive behavior through a fibroblast-associated epithelial program that is molecularly distinct from canonical EMT.

The spatial analysis of the peritoneal metastasis identified a tumor–stroma organization resembling that observed in primary tumors. ECMr and epithelial fractions exhibited widespread spatial coupling, whereas ECMi fibroblasts occupied spatially distinct regions. LAMC2 expression and hemidesmosome-associated gene-program activity were again inversely organized with proliferation, recapitulating the organization observed at the primary tumor interface. These findings show that the ECMr-associated spatial niche and related epithelial program can also be observed at a peritoneal metastatic site. Nevertheless, because the metastatic analysis was based on a single spatial specimen, it should be viewed as proof of principle rather than evidence of a universally conserved metastatic architecture.

From a translational perspective, ECMr-associated GLIS2 regulon activity may provide a means to identify tumors with increased ECMr-associated stromal activity using bulk expression data. Because activity was measured in resected primary tumors, its reproducible association with shorter peritoneal metastasis-free survival across two independent gastric cancer cohorts suggests potential value for biologically stratifying tumors with a greater propensity for subsequent peritoneal dissemination. However, the current analysis does not establish clinical utility, a validated threshold, or superiority over existing clinicopathologic models. Development of a clinically applicable assay will require validation using protein-level markers, targeted expression panels, or spatial pathology approaches in prospective cohorts. In addition, because GLIS2 regulon activity is an indirect surrogate, future studies should determine whether combinations of ECMr-associated markers provide more specific and robust detection of this stromal state.

Several limitations should be considered. First, the inferred fibroblast trajectory was derived from computational analyses of cross-sectional transcriptomic data. Although pseudotime, CytoTRACE2, regulon activity, and spatial gradients were concordant, direct lineage tracing was not performed. Second, higher GLIS2 regulon activity was associated with the ECMr state, but the functional role of GLIS2 was not experimentally validated. Genetic perturbation will be required to determine whether GLIS2 contributes to fibroblast state transition or to the effects of ECMr fibroblasts on cancer cells. Third, the hemidesmosome-associated program was defined at the transcriptomic level, and the inferred reciprocal collagen–LM332 axis was not functionally perturbed. Protein-level localization, ultrastructural confirmation, and perturbation of LM332, α6β4 integrin, or fibroblast-derived collagen will be necessary to establish whether functional hemidesmosome-like adhesions form and contribute to tumor-cell movement. Fourth, although multiple ECMr fibroblast clones were examined, they originated from a single immortalized patient-derived fibroblast preparation and were tested with one gastric cancer cell line. Moreover, the longitudinal imaging was qualitative rather than a formal motility assay. Validation across independent fibroblast donors and cancer models, together with quantitative migration analyses, will therefore be needed. Fifth, the number of spatially profiled primary and metastatic specimens was limited, and the peritoneal metastasis analysis was based on a single case. Validation in larger spatial cohorts is therefore needed. Sixth, the resolution of the Visium platform does not permit unambiguous assignment of the measured signals to individual cells or direct demonstration of cell–cell contact at the tumor–stroma interface. Finally, the clinical analyses relied on GLIS2 regulon activity as a bulk-level surrogate for ECMr-associated stromal activity and did not evaluate prospective prediction or clinical decision benefit.

In summary, this study identifies a spatially organized continuum of ECM fibroblast states in gastric cancer and defines GLIS2 regulon activity as a transcriptional feature of the tissue-remodeling ECMr state. ECMr-associated GLIS2 regulon activity was independently associated with shorter peritoneal metastasis-free survival in two gastric cancer cohorts, with a similar association observed in one colorectal cancer cohort. Interaction inference and co-culture analyses linked this stromal context to a non-EMT epithelial remodeling program characterized by coordinated LM332-subunit expression, a hemidesmosome-associated gene program, a matrix-remodeling protease profile associated with wound re-epithelialization, and reduced proliferative activity, while longitudinal imaging showed progressive elongation and positional displacement of cancer cells over the ECMr fibroblast layer. Related stromal and epithelial organization was observed at the primary tumor–stroma interface and in a peritoneal metastasis. Together, these findings define an ECMr-centered stromal–epithelial wound-repair circuit that couples fibroblast remodeling to an LM332–hemidesmosome-associated, low-proliferative epithelial state, supports epithelial movement at the tumor–stroma interface, and is linked to a heightened propensity for subsequent peritoneal dissemination.

## METHODS

### Human participants

Spatial transcriptomic profiling was performed on two primary gastric cancer sections (ST1, ST2), one non-neoplastic gastric section (NT1), and one peritoneal metastasis section (PT1). ST1 and ST2 were obtained from two patients with SFRP4-high gastric cancer defined by bulk expression measurement using the nProfiler 1 Stomach Cancer Assay (Novomics, Seoul, Republic of Korea), a clinically validated RT-qPCR-based in vitro diagnostic kit. These patients underwent curative gastrectomy at Yonsei University Hospital, with tissue blocks sampled at the tumor invasive edge to capture the tumor-stroma interface (IRB No. 1-2020-0004). NT1 was obtained from the distal resection margin of an independent gastrectomy specimen at Seoul National University Hospital and processed as a histologically intact reference (IRB No. H-2106-156-1230). PT1 was reanalyzed from a publicly available dataset generated by our group as part of the sampling cohort described in Lee et al.^31^, in which the PT1 specimen itself was not reported. No new tissue acquisition was performed for PT1. Clinicopathologic characteristics and bulk gene expression values for the four marker genes (SFRP4, *GZMB*, *WARS*, and *CDX1*) of the spatial transcriptomic samples are summarized in Table S1. Patient-derived fibroblasts (pdFib) used for in vitro validation were isolated from anonymized gastric cancer tissue obtained at Yonsei University Hospital, from which four monoclonal lines were subsequently established (IRB No. 4-2022-0791). Detailed donor clinicopathologic information was not retained in accordance with the anonymization protocol. Informed consent was obtained from all participants prior to tissue collection.

### Cell lines

The SNU668 human gastric cancer cell line was obtained from the Korean Cell Line Bank. HEK293T cells were used for lentivirus production. Immortalized patient-derived fibroblast monoclonal lines (C1-C4) and the parental immortalized pool were generated in this study as described below. The immortalized monoclonal lines were maintained in DMEM supplemented with 10% fetal bovine serum (FBS), 100 ug/mL Primocin, and 1% penicillin-streptomycin at 37 °C with 5% CO2. All cell lines were confirmed to be negative for mycoplasma contamination.

### Spatial transcriptomics sample collection and processing

ST1 and ST2 tissue blocks were sampled at the tumor invasive edge to capture the tumor-stroma interface, and NT1 was processed from the distal resection margin of an independent gastrectomy specimen as a histologically intact reference. Spatial transcriptomics was performed using the 10x Genomics Visium Spatial Gene Expression v1 platform on formalin-fixed paraffin-embedded gastric tissue sections (6.5 x 6.5 mm) stained with hematoxylin and eosin. Library preparation and sequencing for ST1 and ST2 were conducted by Geninus Inc. (Seoul, Korea), and raw reads were processed using Space Ranger (v1.1.0). Library preparation and sequencing for NT1 were conducted by Macrogen Inc. (Seoul, Korea) and raw reads were processed using Space Ranger (v1.3.0). Both pipelines used the GRCh38-2020-A reference genome. PT1 was reanalyzed from previously published count matrices.

### Single-cell RNA-seq processing and cell type annotation

Publicly available scRNA-seq data of gastric cancer were obtained from the Genome Sequence Archive (HRA003647). Cells were filtered by median absolute deviation (MAD)-based outlier detection, with thresholds of 5 MADs for total counts, detected genes, and the percentage of counts in the top 20 genes, and 3 MADs or an absolute threshold of 20% for mitochondrial gene percentage. Ambient RNA contamination was corrected using SoupX, and genes expressed in fewer than 20 cells were excluded. Expression values were normalized using median-based size factors followed by log1p transformation, and highly variable genes were selected based on binomial deviance. Batch correction was performed using scVI with a 15-dimensional latent space and sample as the batch key, followed by semi-supervised refinement with scANVI. UMAP embedding was computed with k = 50 neighbors and min_dist = 0.1, and clustering was performed using the Leiden algorithm. Cell types were annotated based on canonical marker gene expression, including T cells (*PTPRC*, *CD8A*, *CD4*), B cells (*MS4A1*), myeloid cells (*CD163*, *CXCL8*), mast cells (*KIT*), neutrophils (*CXCL8*), plasma cells (*TNFRSF17*), endothelial cells (*PLVAP*), epithelial cells (*EPCAM*, *CDH1*), and stromal cells (*THY1*, *COL1A1*). Clusters with mixed or undefined marker expression were excluded from downstream analysis.

### Epithelial cell clustering and malignant cell identification

Epithelial cells annotated from the major cell type clustering were extracted for sub-clustering analysis. SoupX-corrected counts were normalized by total count followed by log1p transformation, and the top 3,000 highly variable genes were selected. The expression matrix was scaled (max value = 10) and reduced by PCA, and a neighborhood graph (k = 50, 10 principal components) was constructed for UMAP embedding (min_dist = 0.01) and Leiden clustering (resolution = 1.0). Sub-clusters were annotated as pit (Pit), chief (Chief), isthmus (Isthmus), and malignant (Malig) cells based on canonical marker gene expression (*TFF1*, *GKN1*, *MUC5AC* for pit cells; *LIPF*, *MUC6* for chief cells; *STMN1* for isthmus cells) and inferCNV-based copy number alteration signal^55^. Genome-wide copy number alterations were inferred with inferCNV using normal-origin sub-clusters as reference, with cutoff 0.1, denoising (noise filter 0.12), subcluster mode, i6 HMM model, and Ward.D2 hierarchical clustering. Malignant cells were identified as tumor-origin sub-clusters with elevated genome-wide CNV burden relative to the normal-origin reference.

### Fibroblast subtype identification and trajectory analysis

The stromal subset was re-processed independently with an additional mitochondrial gene percentage filter (<10%). Highly variable genes were selected using DUBStepR, and batch correction was performed using scVI with a 10-dimensional latent space and sample as the batch key, followed by scANVI refinement. Leiden clustering at resolution 0.8 identified six fibroblast subtypes and two mural cell populations. Fibroblast subtypes were annotated as MMP1+ (*MMP1*, *COL7A1*, *ISG15*), SOX6+ (*SOX6*, *NRG1*, *POSTN*), APOE+ (*CTSC*, *CCL11*, *APOE*), ECMh (*MFAP5*, *CD34*, *CLEC3B*), ECMi (*C7*, *SFRP2*, *CXCL12*), and ECMr (*COL8A1*, *TIMP1*, *FN1*), with shared ECM lineage markers including *OGN*, *DCN*, *LUM*, and *CCDC80*. Mural cell populations comprised smooth muscle cells (*DES*, *ACTG2*, *ACTA2*) and pericytes (*CSPG4*, *RGS5*, *NOTCH3*, *MCAM*). Developmental relationships among stromal populations were inferred using partition-based graph abstraction (PAGA) on a force-directed layout initialized from the UMAP embedding, with diffusion pseudotime (DPT) calculated using ECMh as the root state. Cellular potency scores were estimated using CytoTRACE2 on the raw count matrix (full model, batch size = 1,000, maximum 200 principal components, random seed = 1) to assign per-cell developmental potential scores and categorical potency labels (differentiated, unipotent, oligopotent, multipotent).

For differential pathway activity between ECMi and ECMr fibroblasts, per-cell scores were inferred with a univariate linear model (ULM, decoupleR) on curated MSigDB Hallmark and GO Biological Process gene sets, and t-scores were calculated by Welch’s t test (ECMr versus ECMi).

### Regulon analysis

Transcription factor regulon activities were inferred using the pySCENIC pipeline (v0.12.1). Gene regulatory networks were inferred with GRNBoost2 from the log-normalized expression matrix using a curated human TF list. Regulon prediction and motif enrichment were performed using two cisTarget ranking databases for hg38 (10 kb upstream/downstream and 500 bp upstream/100 bp downstream of the TSS) with the motifs-v9-nr annotation table. Per-cell regulon activity scores were quantified using AUCell. Branched expression analysis was performed using CytoTRACE2 relative potency scores as the ordering variable, with ECMi treated as the shared branch point. The ECMh–ECMi arm was displayed in reverse order to represent the proposed ECMh-to-ECMi transition (Transition I), followed by the ECMi-to-ECMr arm (Transition II). For each regulon, generalized additive models (GAMs) were fitted independently along each branch, and branch divergence was quantified as the mean absolute error between fitted curves over the shared distance range, normalized by the pooled standard deviation. Statistical significance was assessed by a permutation test (1,000 permutations) shuffling branch labels of non-ECMi cells while preserving ECMi as the shared branch point, with Benjamini-Hochberg correction. Significant regulons were hierarchically clustered (Ward’s D2) on z-scored smoothed expression profiles, and Pearson correlations between regulon activities and CytoTRACE2 scores were computed separately for each phase.

### Spatial transcriptomics deconvolution

Cell type abundances at each spot were estimated using Cell2location. A reference signature matrix was constructed from the scRNA-seq dataset (HRA003647) using the regression model with all annotated fibroblast and epithelial subtypes, with sample as the batch key (500 epochs, 1,000 posterior samples). Genes were filtered by cell count (>=10), cell percentage (>3%), and non-zero mean expression (>1.12). For spatial mapping, mitochondrial genes were removed prior to deconvolution. The mapping model was trained on each tissue section independently with expected cell abundance per location set to 5 and detection alpha set to 20 (10,000 epochs), and The 5th percentile of the posterior cell-abundance distribution (q05) was used as a conservative estimate of cell-type abundance. For downstream analyses, q05 estimates across the 17 mapped cell populations were row-normalized within each spot to sum to 1 and are reported as inferred cell-type fractions.

### Epithelial-stromal boundary analysis

To localize the epithelial-stromal interface, a gradient-based boundary score was computed from Cell2location fraction maps. Cell type abundances were first normalized to row-sum fractions per spot, and a spatial neighbor graph was constructed using six nearest neighbors (hexagonal Visium grid). For each spot, spatial gradients of epithelial and stromal fractions were estimated from neighboring spots, and the boundary score was defined as the product of gradient magnitudes weighted by the inverse of cosine similarity between gradient vectors, such that anti-directional gradients (one fraction increasing while the other decreases) received higher scores. Spots above the 75th percentile of the boundary score distribution were designated as the boundary zone. Within the boundary zone, abundances of ECMh, ECMi, and ECMr were compared pairwise using the Mann-Whitney U test to identify the dominant ECM fibroblast subtype at the epithelial-stromal interface.

### Spatial niche identification

Spatial domains were identified using a Local Indicators of Spatial Association (LISA)-based clustering approach applied to Cell2location-derived cell type composition maps. Cell type abundances were normalized to row-sum fractions across four major categories (epithelial, stromal, immune, smooth muscle). A Gaussian kernel-based spatial weight matrix was constructed from spot coordinates (k = 10 neighbors), and univariate Local Moran’s I statistics were computed for each major cell type with 999 permutations. Only spots in the High-High (HH) quadrant were retained. Cell type-specific noise filtering was applied using quantile-based thresholds on raw abundance and LISA values, with differential scaling factors (immune scale = 3.0, stromal scale = 1.0). Filtered LISA features were processed through a dual scaling strategy combining feature-wise z-score standardization and sample-wise relative abundance normalization. The combined feature matrix was reduced by PCA, followed by shared nearest neighbor graph construction and Leiden clustering (resolution = 0.8). Final niche annotations (epithelial, stromal, immune, smooth muscle) were assigned by inspecting LISA feature enrichment patterns per cluster.

### Spatial distance and trajectory analysis

To quantify the spatial proximity between stromal and malignant compartments, a stepwise tier distance metric was computed based on the hexagonal grid structure of the Visium platform. For each stromal niche spot, the minimum number of hexagonal grid steps required to reach the nearest epithelial niche spot was determined using a breadth-first search algorithm over the spatial neighbor graph. This tier distance was applied in multiple contexts throughout the study, including distance from stromal to epithelial niches (“distance to cancer”), distance from epithelial to stromal niches (“distance to stromal”), and distance from tissue spots to the peritoneal surface (“distance to peritoneal surface”). To assess the spatial relationship between ECMi and ECMr fibroblast subtypes within the stromal niche, bivariate Moran’s I was computed using the PySAL library on z-score standardized fractions, with a K-nearest neighbor spatial weight matrix (k = 6, row-normalized) and 999 permutations. Local bivariate Moran’s I was additionally computed to classify spots into High-High, Low-High, High-Low, and Low-Low quadrants (p < 0.05).

To spatially resolve the ECMi-to-ECMr transition within the stromal niche, spatial trajectory analysis was performed on stromal-assigned spots. Spatially variable genes were identified using Moran’s I autocorrelation (squidpy, FDR-corrected p < 1e-10, I > 0.1). The expression matrix was scaled (max value = 10) and reduced by PCA (50 components), and a neighborhood graph (k = 50) was constructed for UMAP embedding and Leiden clustering (resolution = 1.0). Diffusion maps were computed from the PCA representation, the diffusion map-based neighbor graph was used for PAGA connectivity estimation, and diffusion pseudotime was calculated with the cluster exhibiting the highest ECMi fraction designated as the root state. The inferred pseudotime was then mapped onto tissue spatial coordinates. Per-spot transcription factor regulon activities were estimated by converting SCENIC-derived regulon target gene sets into a decoupleR-compatible network (retaining the maximum weight for duplicate TF-target pairs) and applying the univariate linear model (ULM) method to the log-normalized Visium expression matrix, with Pearson correlation between regulon activity and tier distance used to identify spatially graded regulons.

### Cell-cell communication analysis

Intercellular communication networks were inferred using CellChat (v1.6.1) with the full CellChatDB human database, encompassing secreted signaling, ECM-receptor, cell-cell contact interactions, and non-protein interaction. For the scRNA-seq dataset, communication probabilities were computed from the log-normalized count matrix using the default mass action-based model with cell type annotations as group identifiers, and interactions involving fewer than 50 cells in either the sender or receiver group were filtered out. Aggregated communication networks were constructed by summing interaction counts and weights across all significant ligand-receptor pairs, and network centrality scores were computed to identify dominant sender and receiver roles. For spatial transcriptomics data, CellChat was applied with distance-aware communication probability estimation. Spots were classified into proximal and distal categories based on the tier distance metric, yielding six niche groups (Proximal_Stromal, Distal_Stromal, Proximal_Malig, Distal_Malig, Immune, SMC). Tissue coordinates and scale factors were extracted from the Space Ranger output to convert pixel distances to physical units (theoretical spot size = 65 um). Communication probabilities were computed using a truncated mean approach (trim = 0.1) with distance-dependent decay (interaction range = 250 um, scale factor = 0.01) and contact-dependent signaling (contact range = 100 um), and interactions involving fewer than 10 spots were filtered out.

### Patient-derived fibroblast isolation and immortalization

Patient-derived fibroblasts (pdFib) were isolated from anonymized gastric cancer tissue obtained at Yonsei University Hospital. Freshly resected tissues were transferred to culture medium on ice, washed in phosphate-buffered saline containing 500 U/mL penicillin-streptomycin, and minced into approximately 1 mm3 fragments using scalpels. Tissue fragments were plated in 10 cm dishes with 7 mL of culture medium and incubated at 37 C with 5% CO2 and 95% humidity. Fibroblast outgrowth was typically observed within 5 days, and isolated fibroblasts were used between passages 4 and 6. pdFibs were immortalized by lentiviral transduction of SV40 large T antigen with a tdTomato reporter (pLenti-EF1a-LTA-IRES-tdTomato). Following transduction, monoclonal populations were isolated by fluorescence-activated cell sorting (FACS) using a BD FACS Aria II SORP Cell Sorter (BD Biosciences), and four monoclonal lines were established by clonal expansion for 4 weeks in 96-well plates followed by an additional 3 weeks before transfer to 100 mm dishes. SV40 large T antigen expression in the established clones was confirmed by RT-qPCR with expression levels comparable to HEK293T. The immortalized monoclonal lines were maintained in DMEM (Gibco) supplemented with 10% fetal bovine serum (FBS, Gibco), 100 ug/mL Primocin (InvivoGen), and 1% penicillin-streptomycin (Gibco) at 37 C with 5% CO2.

### Plasmid construction and lentivirus production

pLenti-EF1a-LTA-IRES-tdTomato was generated by Gibson assembly of three fragments: an AgeI-digested pLenti-PE2max-BSD backbone (Addgene, 191102), SV40 large T antigen amplified from pBABE-puro SV40 LT (Addgene, 13970), and tdTomato amplified from pLVX-IRES-tdTomato-FlagAkt1 (Addgene, 64831). pLenti-EF1a-eGFP was generated by Gibson assembly of an FseI/MluI-digested Lenti_gRNA-Puro backbone (Addgene, 84752), the EF1a promoter, and eGFP amplified from pEGFP (Addgene, 165830). All primers are listed in Table S9, and oligonucleotides were synthesized by Macrogen. For lentivirus production, HEK293T cells were seeded at 5 x 10^6 cells per 100 mm dish 24 hours before transfection. Medium was replaced with DMEM containing 25 uM chloroquine diphosphate for 5 hours. Transfer plasmid, psPAX2, and pMD2.G were mixed at a molar ratio of 1.64, 1.3, and 0.72 pmol, respectively, diluted in 500 uL Opti-MEM (Life Technologies), and complexed with polyethylenimine (PEI) at a 1:3 DNA-to-PEI weight ratio for 20 minutes before addition to cells. Caffeine (Sigma-Aldrich) was added at 4 mM final concentration to enhance viral titer. At 12 hours post-transfection, medium was refreshed with 4 mM caffeine-supplemented growth medium. Viral supernatant was harvested at 36 hours, centrifuged at 2,000 x g for 10 minutes, filtered through 0.45 um low protein-binding membranes (Millipore), and stored at -80 C.

### RNA extraction and bulk RNA-seq processing

Total RNA was extracted from up to 2 x 10^6 cells using the TRIzol method, and 800-1,000 ng of RNA was used for library preparation with the Illumina TruSeq RNA Sample Prep Kit. Libraries were sequenced in paired-end mode at Macrogen (Seoul, Korea) on either the Illumina NovaSeq 6000 or NovaSeq X Plus platform. Raw FASTQ files were processed through the TOPMed/GTEx harmonized RNA-seq pipeline. Reads were aligned to the GRCh38 reference genome using STAR (two-pass mode) with parameters specified by the TOPMed pipeline, including --outFilterMultimapNmax 20, --alignSJoverhangMin 8, --alignSJDBoverhangMin 1, -- outFilterMismatchNoverLmax 0.1, --alignIntronMin 20, --alignIntronMax 1000000, --quantMode TranscriptomeSAM GeneCounts, and --chimSegmentMin 15. Transcriptome-aligned BAM files were then quantified using RSEM (rsem-calculate-expression) with --paired-end --estimate-rspd --fragment-length-max 1000, producing per-gene expected counts and TPM values used for downstream analyses.

### ECM fibroblast subtype identification of immortalized fibroblasts

Prior to co-culture, the four immortalized monoclonal pdFib lines were validated as the ECMr subtype using bulk RNA-seq profiling, with the parental immortalized pool serving as the reference. Genes were filtered by minimum expression (log2 TPM >= 0.5 in at least one sample). For ECM subtype scoring, signature gene sets for ECMh, ECMi, and ECMr were derived from the scRNA-seq fibroblast cluster markers identified by Seurat FindAllMarkers, retaining genes with adjusted p-value < 1e-30 and average log2 fold change > 1 per cluster ranked by average log2 fold change. Per-sample signature scores were computed using the scanpy gene set scoring function on log2 TPM-normalized expression. For GLIS2 regulon activity, the SCENIC regulon output was converted into a decoupleR-compatible network by extracting TF-target-weight triplets from the TargetGenes field and aggregating duplicate TF-target pairs by retaining the maximum weight. Per-sample regulon activity scores were then estimated using the decoupleR ULM method on log2 TPM-normalized expression.

### Tumor-fibroblast co-culture and transcriptomic profiling

For transcriptomic profiling of cancer cells exposed to ECMr fibroblasts, immortalized monoclonal pdFibs and SNU668 gastric cancer cells (Korean Cell Line Bank) were directly co-cultured. Prior to co-culture, SNU668 cells were transduced with pLenti-EF1a-eGFP and pdFibs with pLenti-EF1a-LTA-IRES-tdTomato, and the labeled populations were isolated by FACS after 14 days. The two cell types were seeded at a 1:3 (pdFib:SNU668) ratio in 100 mm dishes with DMEM supplemented with 10% FBS, 100 ug/mL Primocin, and 1% penicillin-streptomycin, maintained for 14 days with medium changes every 2 days, and transferred to 150 mm dishes upon reaching 80% confluency. Time-matched monoculture controls consisted of GFP-labeled SNU668 cells maintained in parallel for 14 days under the same culture conditions without fibroblasts. Following co-culture, GFP+ SNU668 cells were isolated by FACS, and post-sort purity was assessed by re-gating the sorted samples to quantify residual fibroblast contamination. Five independent co-culture samples generated using C2 (n = 1), C3 (n = 2), and C4 (n = 2) were retained for downstream transcriptomic analysis. Bulk RNA-seq libraries were prepared, sequenced, and processed as described in the RNA extraction and bulk RNA-seq processing section. For differential expression analysis, raw counts were filtered using decoupleR (min_count = 10, min_total_count = 15, large_n = 10, min_prop = 0.7), and PyDESeq2 was applied with group as the design factor. The Wald test was used to contrast co-cultured cancer cells against 14-day time-matched monoculture controls, with Benjamini-Hochberg correction. Pathway-level functional characterization was performed by applying the decoupleR ULM method to the Wald statistic vector against three reference networks independently: MSigDB Hallmark (v7.4) and Gene Ontology Biological Process gene sets (c5.go.bp.v7.4) for pathway activity, CollecTRI for transcription factor activity, and the human PROGENy model for signaling pathway activity, with significance defined at FDR < 0.05. Cell cycle-associated differentially expressed genes (adjusted p < 0.05) were separately extracted using the cell cycle gene list reported by Tirosh et al.^56^. to define the “proliferation signature” used in subsequent spatial analyses. For visualization of individual gene expression or pathway activity scores in bar plots comparing WT and co-culture groups, Welch’s t-test was used to assess statistical significance.

### Longitudinal imaging of cancer cells on fibroblast monolayers

To monitor the cellular behavior of cancer cells on ECMr fibroblasts over time, immortalized monoclone 2 was seeded at 1.5 x 10^5 cells per well in 24-well plates. Fiducial markers for longitudinal tracking were established by marking random regions of interest (ROI) on the plate surface prior to cell seeding. The following day, fibroblasts were treated with mitomycin C (10 ug/mL) for 2 hours to arrest proliferation, after which the medium was replaced. SNU668-GFP cancer cells were seeded at 3 x 10^4 cells per well 24 hours after mitomycin C treatment. Beginning from the day after cancer cell seeding (designated Day 0), daily imaging was performed by first capturing the fiducial markers under brightfield to ensure consistent field alignment across timepoints, followed by GFP fluorescence image acquisition at identical positions. Monoculture controls without fibroblasts were prepared in parallel under identical conditions. For image analysis, the green channel was extracted from each acquired image using ImageJ with brightness and contrast standardized across timepoints, and fiducial markers were manually annotated for each timepoint to align images into a common coordinate system. GFP images were smoothed by Gaussian filtering, background-subtracted, and thresholded to extract cell-occupied regions, which were overlaid across all timepoints (Day 0, Day 1, Day 4, Day 7) with timepoint-specific colors. Positional displacement and morphological changes of cancer cells were assessed by direct visual comparison between co-cultured and monoculture conditions.

### Spatial validation of in vitro transcriptional programs

To spatially validate the co-culture-induced transcriptional programs in malignant cells, pathway activity scores for the proliferation signature and hemidesmosome assembly (GOBP) were computed per Visium spot. The proliferation signature was defined from the cell cycle-associated differentially expressed genes derived from the co-culture bulk RNA-seq analysis, and per-spot scores were computed using the scanpy gene set scoring function. Hemidesmosome assembly activity was estimated using the decoupleR ULM method on the log-normalized expression matrix. For the spatial exclusivity analysis, epithelial niche spots were binarized using the 90th percentile thresholds for the hemidesmosome assembly score and LAMC2 expression and a zero threshold for the proliferation signature score, then assigned to four categories (program-high only, proliferation-high only, both high, or neither). Spatial exclusivity was tested by Fisher’s exact test on the resulting 2 x 2 contingency table, and sensitivity analysis across multiple quantile thresholds (70th-90th percentiles) for the hemidesmosome score confirmed robustness of the association. To evaluate the spatial gradient of these programs relative to the tu ㄹ mor-stroma interface, z-scored pathway activity values were plotted as a function of tier distance to the stromal compartment, with 95% confidence intervals computed from the standard error of the mean at each distance.

### External scRNA-seq validation

To validate the ECM fibroblast subtypes, two independent scRNA-seq datasets were analyzed: a publicly available gastric cancer cohort from Kumar et al. and an in-house Korean gastric cancer cohort previously generated by our group. For the Kumar et al. cohort, raw counts were processed with Seurat using log-normalization, DUBStepR-based highly variable gene selection, scaling, PCA, and Harmony batch correction (30 dimensions), followed by UMAP and Leiden clustering (resolution = 0.6). For the in-house Korean cohort, the stromal compartment was extracted from the integrated atlas after removing scDblFinder-classified doublets and cells with mitochondrial gene percentage >=10%. Counts were normalized (log1p with median-based size factors), and highly variable genes were selected from a precomputed DUBStepR list. Batch correction was performed in two sequential rounds: scVI (n_latent = 10) followed by removal of contaminant and pericyte clusters, then retrained scVI (n_latent = 15) with scANVI supervision using the first-round labels, with subsequent UMAP and Leiden clustering (resolution = 1.0). For both cohorts, fibroblast subtypes were annotated using the marker gene panel defined in the primary cohort, and the developmental trajectory and key regulon activities were assessed using the same analytical pipeline.

### Bulk-cohort survival analysis

To evaluate the prognostic significance of GLIS2 regulon activity, three independent bulk transcriptomic cohorts were analyzed: two gastric cancer cohorts (GSE84437, n = 329; ACRG cohort GSE62254, n = 298) and one colorectal cancer cohort (AMC-AJCCII-90, GSE33113, n = 86). Across all cohorts, only patients with complete metastasis status and survival information were included in the analysis. Peritoneal metastasis-free survival (PMFS) was calculated from the date of surgery using the postoperative follow-up time and recurrence-site annotations available for each cohort. In GSE84437, peritoneal recurrence documented at the first recurrence was considered an event, including cases with synchronous peritoneal and hematogenous recurrence. Patients without a recorded peritoneal recurrence were censored according to the last available follow-up information. For ACRG, the postoperative follow-up and peritoneal recurrence annotations provided with the cohort were used. For each cohort, log2 transformed expression matrices were used to compute per-sample GLIS2 regulon activity scores using the decoupleR ULM method on the SCENIC-derived regulon network. Patients were dichotomized into GLIS2-high and GLIS2-low groups using the maximally selected rank statistic (maxstat) procedure, which identifies the optimal cutpoint maximizing the standardized log-rank statistic. For overall survival analysis, Kaplan-Meier curves were generated and compared between groups using the log-rank test. For peritoneal metastasis-free survival, the maxstat procedure was applied with peritoneal metastasis as the event variable. Univariate Cox regression was performed across recurrence types (peritoneal, hematogenous, lymph node, no recurrence) to identify metastasis patterns associated with GLIS2 activity. Multivariable Cox proportional hazards regression was performed adjusting for T stage, N stage, age, and sex to assess the independent prognostic value of GLIS2 activity, with hazard ratios and 95% confidence intervals reported and presented as summary tables. Distributional differences in GLIS2 activity between metastatic patterns were assessed by Welch’s t-test.

### STATISTICAL ANALYSIS

All analyses were performed in Python (v3.10.20). Two-group comparisons of continuous variables were performed using Welch’s t-test or the Mann-Whitney U test depending on distributional assumptions, and categorical associations were evaluated using Fisher’s exact test. Survival probabilities were estimated by the Kaplan-Meier method and compared by log-rank test, with optimal dichotomization cutoff values determined by the maximally selected rank statistic procedure. Multivariable Cox proportional hazards regression was used to evaluate independent prognostic factors. Differential gene expression from bulk RNA-seq count data was performed using PyDESeq2 (v0.5.2) with the Wald test. Spatial autocorrelation was quantified using univariate and bivariate Moran’s I statistics with permutation-based inference (999 permutations). Multiple hypothesis testing was corrected using the Benjamini-Hochberg false discovery rate procedure unless otherwise specified, and statistical significance was defined as p < 0.05 (two-sided) throughout. Key Python packages used include scanpy (v1.11.5), scVI/scANVI (v1.3.3), decoupleR (v2.1.4), lifelines (v0.30.0), PyDESeq2 (v0.5.2), squidpy (v1.6.5), libpysal (v4.13.0), esda (v2.7.0), scipy (v1.15.3), and statsmodels (v0.14.6).

## Data availability

Spatial transcriptomics data of ST1, ST2, and NT1 newly generated in this study have been deposited in the Gene Expression Omnibus under accession GSE330046. Raw sequencing data for both spatial transcriptomics and bulk RNA-seq are available at the Sequence Read Archive under BioProject PRJNA1458466. Publicly available scRNA-seq data of gastric cancer used as the primary analysis cohort are available at the Genome Sequence Archive under accession HRA003647. Spatial transcriptomics data of the peritoneal metastasis sample (PT1) are available as previously published^31^. Bulk transcriptomic cohorts used for survival analysis are available from the Gene Expression Omnibus under accessions GSE84437, GSE62254, and GSE33113. Custom code is available from the lead contact upon reasonable request. Any additional information required to reanalyze the data reported in this paper is available from the lead contact upon request.

## Acknowledgments

We appreciate the support for this research from the Basic Medical Science Facilitation Programme through the Catholic Medical Center of the Catholic University of Korea, funded by the Catholic Education Foundation. In addition, we specially thank Professor Hye Seung Lee (Seoul National University) for generously sharing the normal gastric tissue Visium spatial transcriptomics and single-cell RNA sequencing data, which were invaluable to this study. This study was supported by the National Research Foundation of Korea (NRF) (RS-2024-00454948 and RS-2025-02273009 to T-M.K.) and by the Korea Health Industry Development Institute (KHIDI) and Soonchunhyang University Research Fund, Republic of Korea (RS-2025-02253005 to S.L.).

## Author contributions

Conceptualization: S.L., J.-H.C., T.-M.K.

Methodology: S.L., S.C.

Software: S.L.

Validation: S.L., S.C.

Formal analysis: S.L.

Resources: D.-S.H., J.K., H.H., J.-H.C.

Data curation: S.L.

Writing — original draft: S.L.

Writing — review and editing: S.L., S.C., H.H.K., J.-H.C., T.-M.K.

Visualization: S.L.

Supervision: H.H.K., J.-H.C., T.-M.K.

Project administration: J.-H.C., T.-M.K.

Funding acquisition: S.L., T.-M.K.

## Declaration of generative AI and AI-assisted technologies in the writing process

During the preparation of this work, the authors used Claude (Anthropic) to improve the readability and language of the manuscript. After using this tool, the authors reviewed and edited the content as needed and take full responsibility for the content of the published article.

