## Supplementary Figures for "Spatial profiling identifies a GLIS2-associated fibroblast state linked to peritoneal recurrence in gastric cancer"

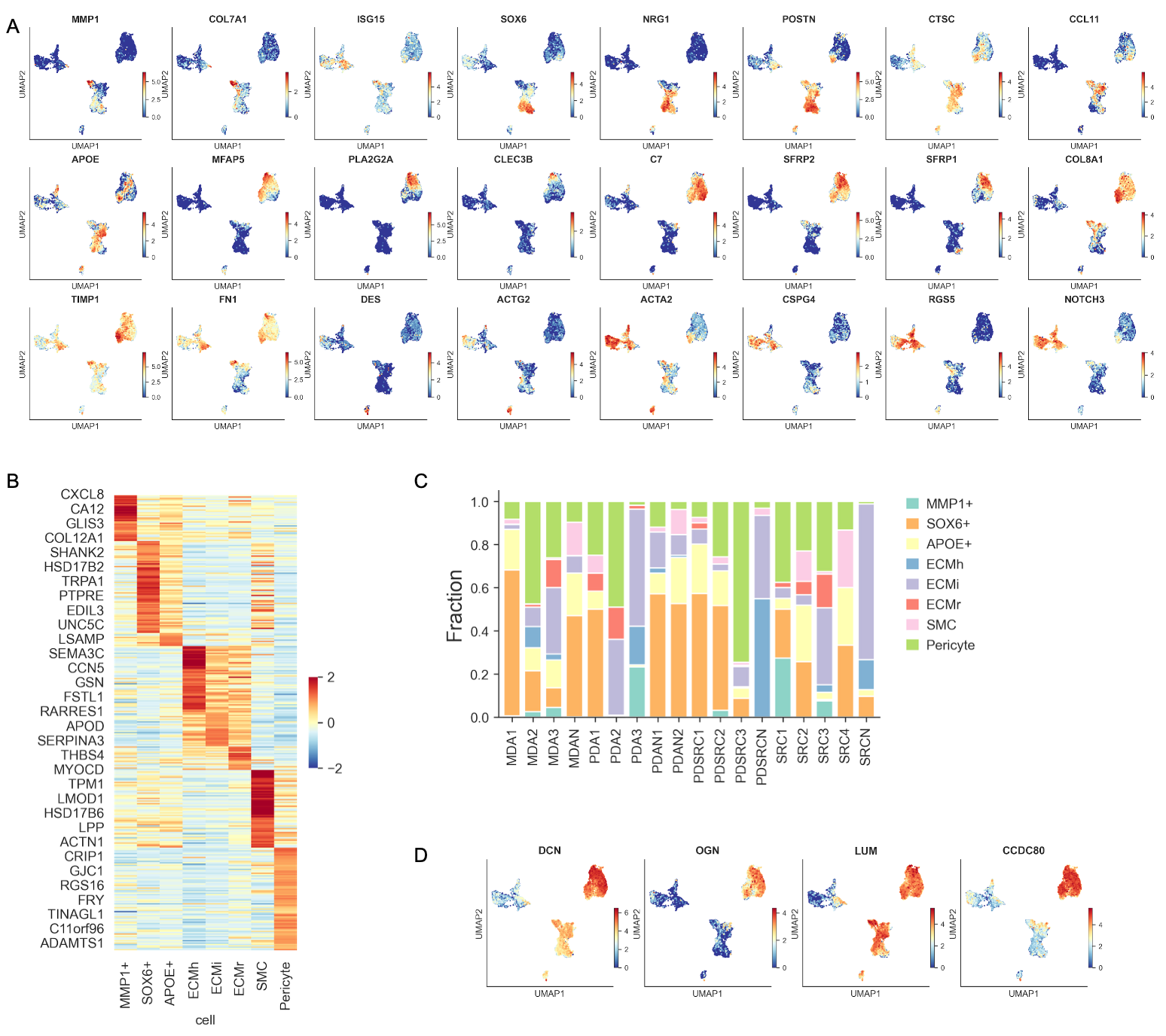

**Figure S1. Identification of stromal cell subsets in gastric cancer scRNA-seq data.** (A) UMAP feature plots of subset-defining marker genes. Color, log-normalized expression. (B) Heatmap of top marker genes across stromal subsets. Values are z-scored expression. (C) Stacked bar plot of stromal subset composition per patient sample. (D) UMAP feature plots of pan-ECM markers (DCN, OGN, LUM, CCDC80).

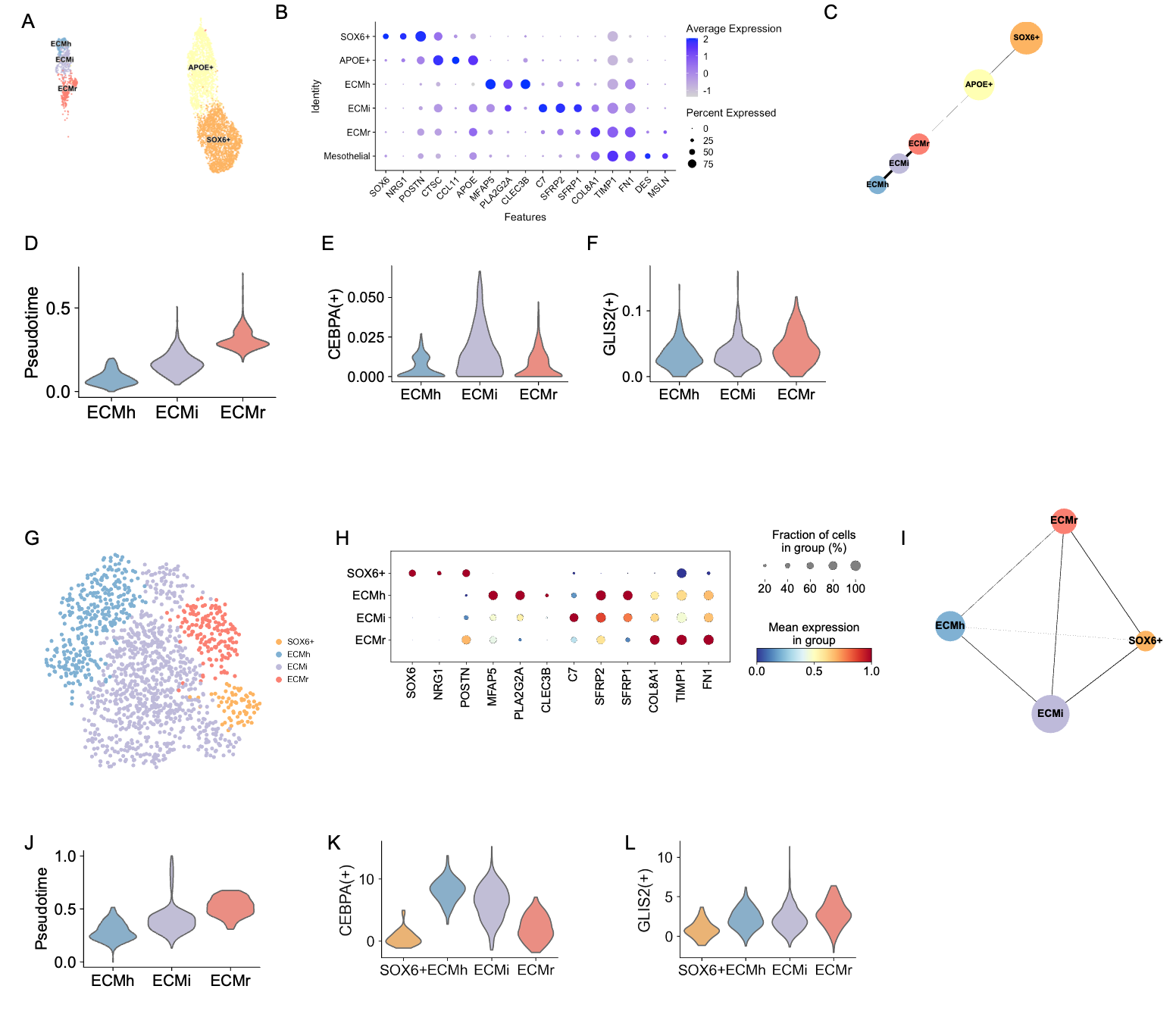

**Figure S2. Reproducibility of ECM fibroblast subsets in independent gastric cancer scRNA-seq cohorts.** (A) UMAP of stromal cells in the Kumar et al. cohort, colored by subset annotation. (B) Dot plot of marker gene expression across stromal subsets. Color, scaled average expression; size, fraction of cells. (C) Partition-based graph abstraction (PAGA) of stromal subsets. Edge thickness, connectivity. (D) Violin plot of diffusion pseudotime in ECMh, ECMi, and ECMr. (E and F) Violin plots of CEBPA(+) and GLIS2(+) regulon activity (AUCell) across ECM fibroblast subsets. (G to L) As in (A to F) for the in-house Korean gastric cancer cohort.

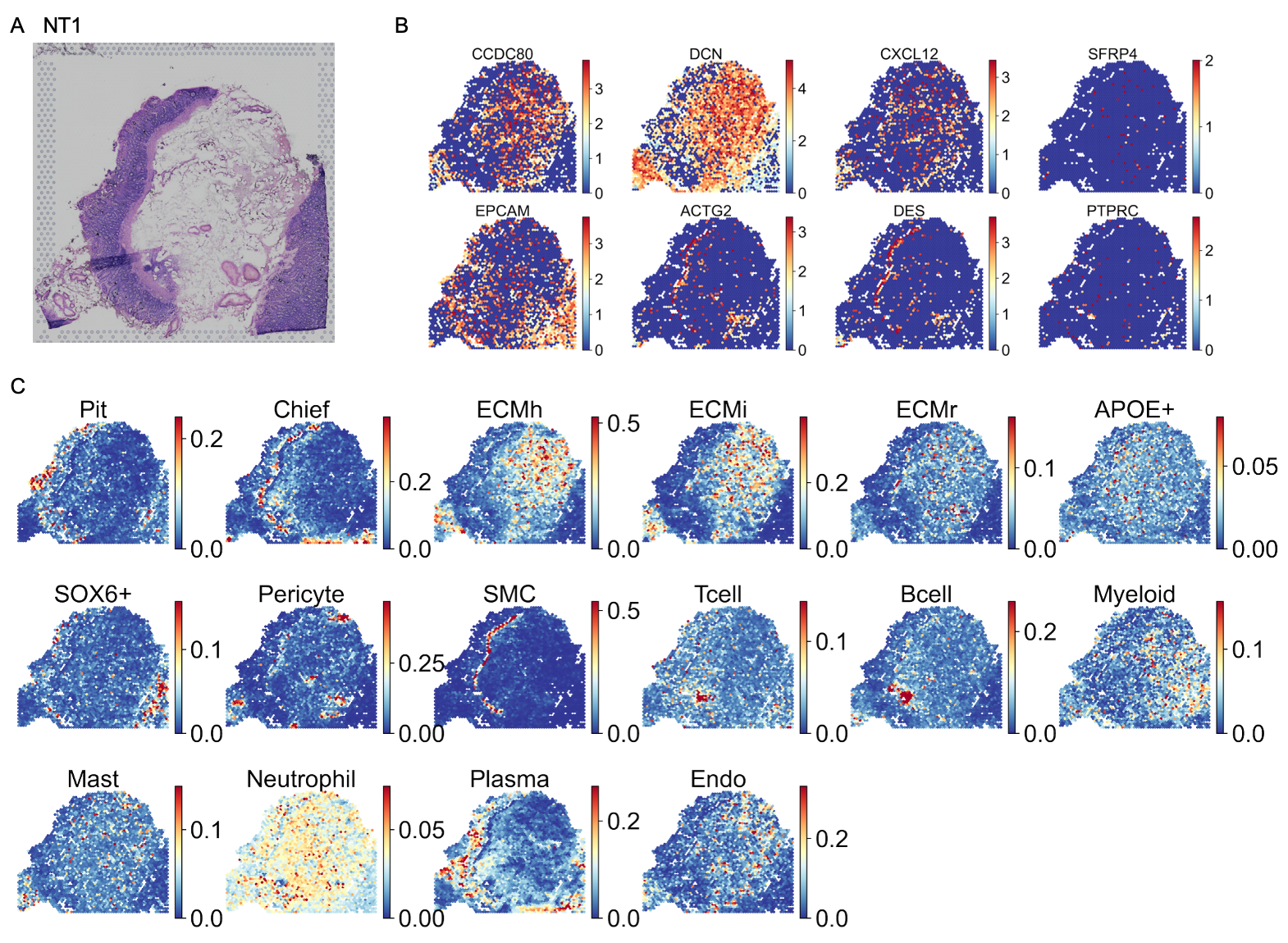

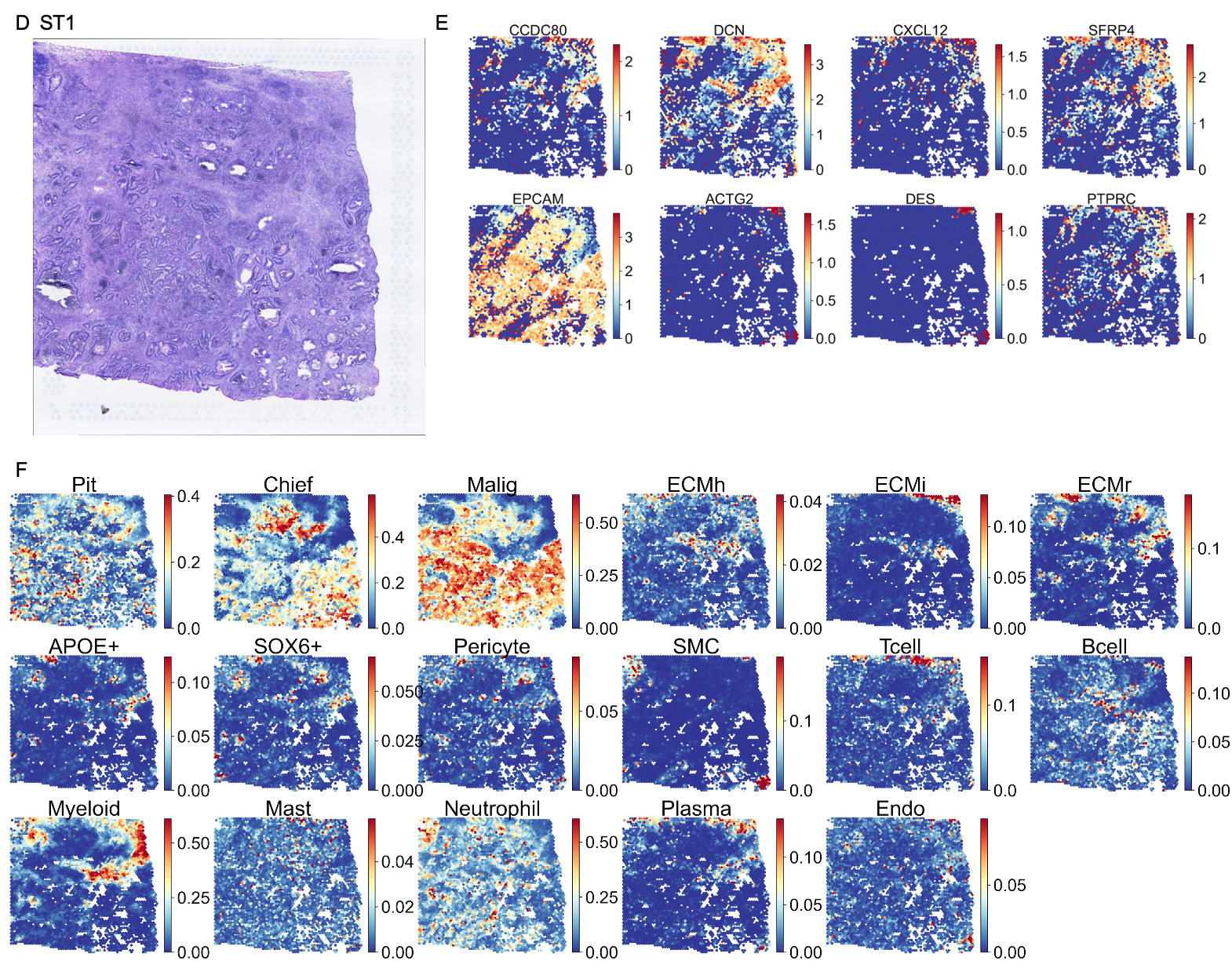

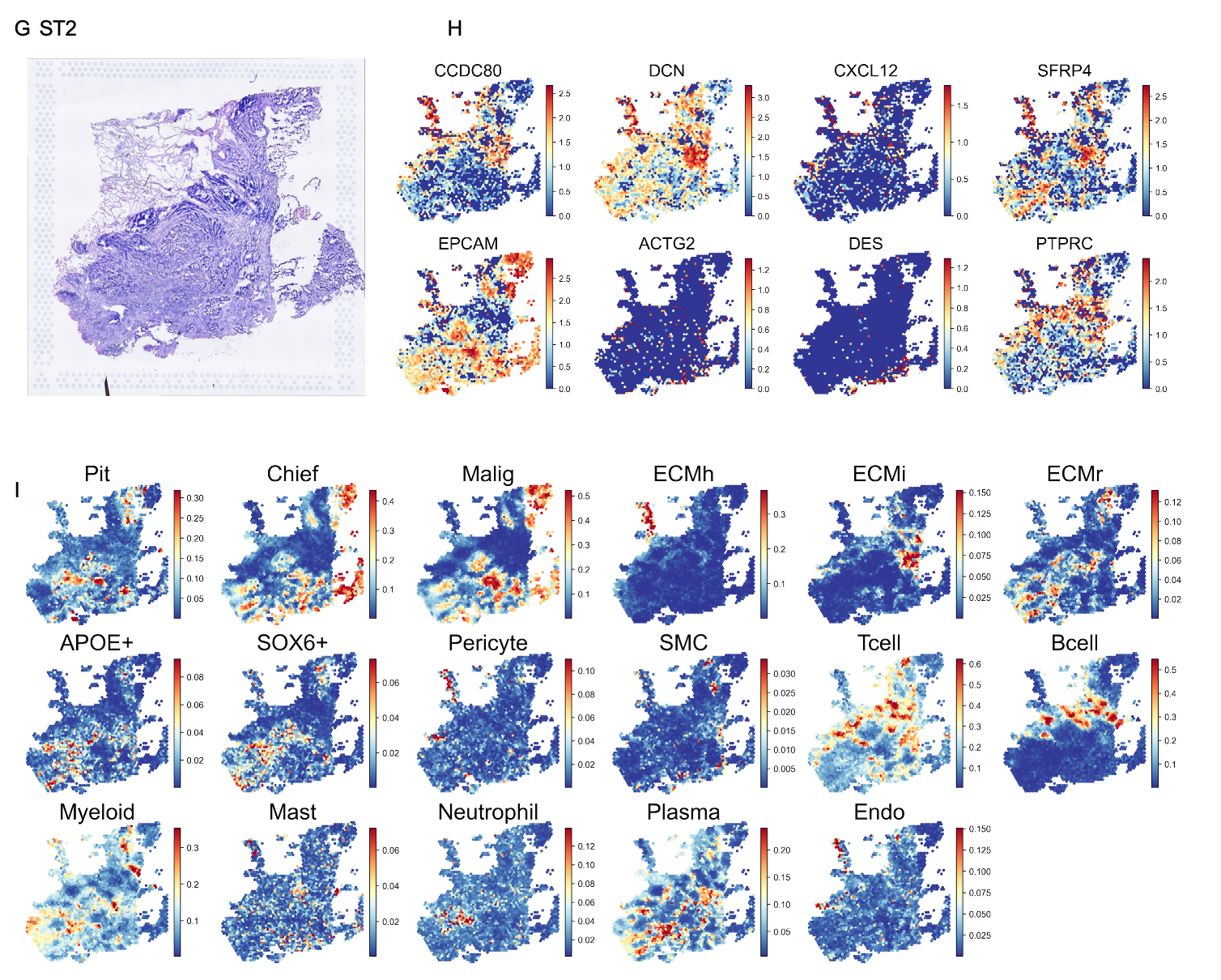

**Figure S3. Histology and spatial cell type maps of NT1, ST1, and ST2.** (A) H&E image of NT1. (B) Spatial expression maps of stromal (CCDC80, DCN, CXCL12, SFRP4), epithelial (EPCAM), smooth muscle (ACTG2, DES), and immune (PTPRC) marker genes in NT1. Color, log-normalized expression. (C) Cell2location-inferred spatial fraction maps of epithelial subsets (Pit, Chief), ECM fibroblasts (ECMh, ECMi, ECMr), other stromal subsets (APOE+, SOX6+, Pericyte, SMC), immune subsets (Tcell, Bcell, Myeloid, Mast, Neutrophil, Plasma), and endothelial cells (Endo) in NT1. Color, estimated cell fraction per spot. (D to F) As in (A to C) for ST1. (G to I) As in (A to C) for ST2.

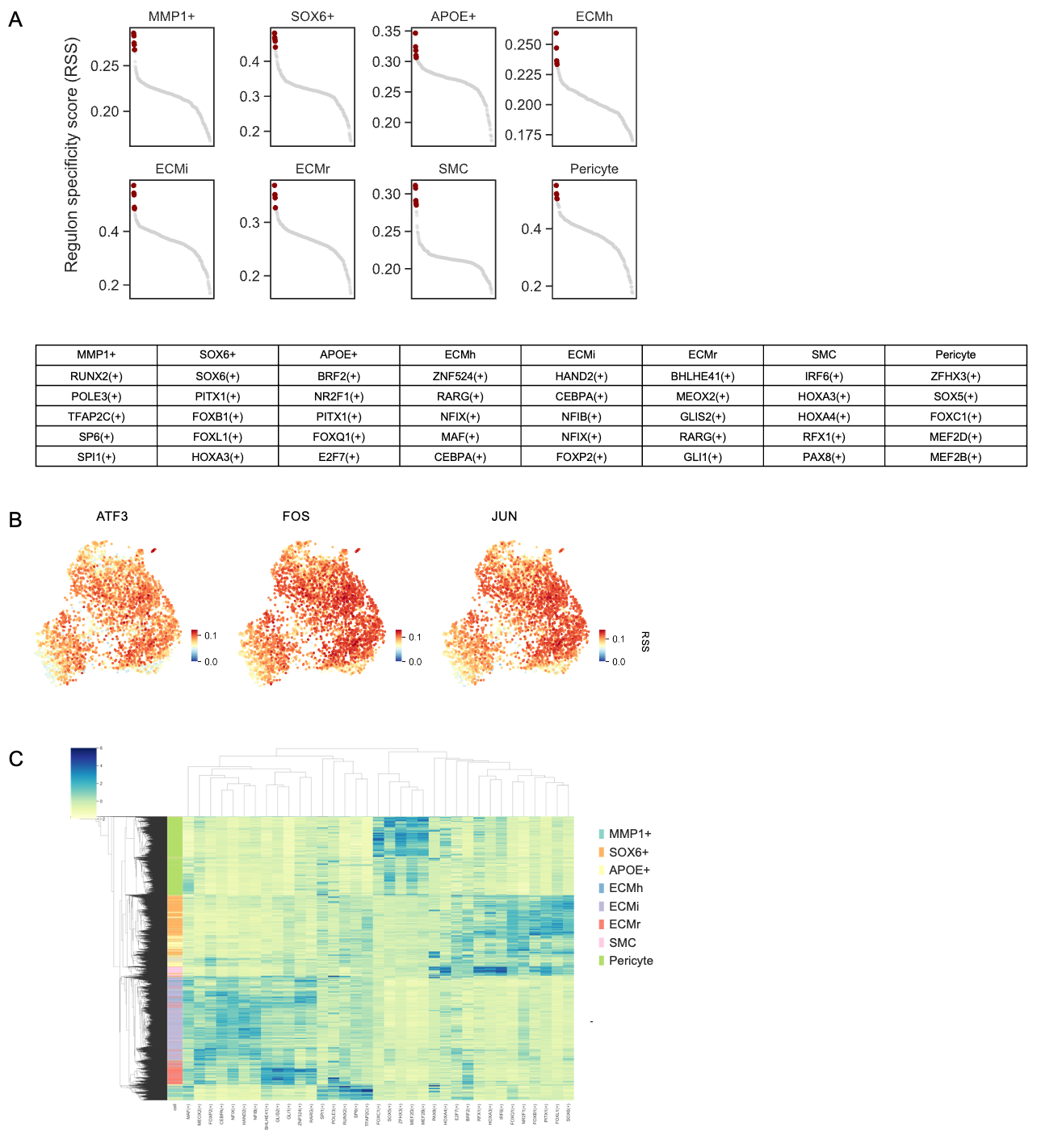

**Figure S4. SCENIC regulon analysis across stromal populations.** (A) Ranked regulon specificity score (RSS) of SCENIC regulons for each stromal subset, with the top 5 regulons highlighted (red) and listed in the table below. (B) UMAP of ECM fibroblast clusters colored by ATF3, FOS, and JUN regulon activity (AUCell). (C) Hierarchically clustered heatmap of regulon activity across stromal subsets. Values are z-scored AUCell activities. Side bar, subset annotation.

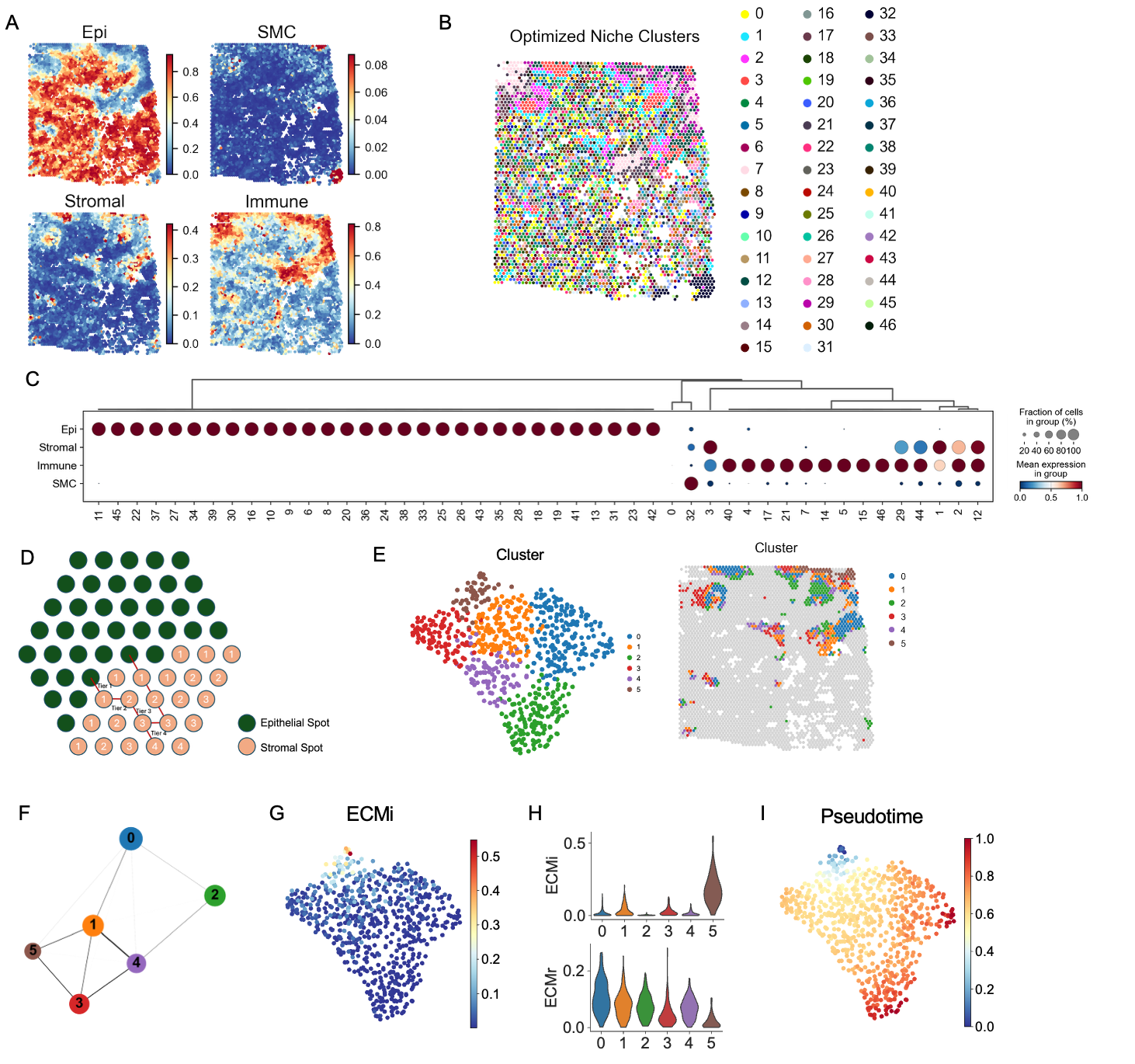

**Figure S5. Spatial niche definition and stromal trajectory analysis in ST1.** (A) Spatial maps of cell2location-inferred Epi, SMC, Stromal, and Immune fractions in ST1, aggregated by major cell type. (B) Spatial map of optimized niche clusters from Local Indicators of Spatial Association (LISA) clustering. (C) Dot plot of compartment marker expression across niche clusters. Hierarchical clustering identifies four major niches (Epi, Stromal, Immune, SMC). Dot size, fraction of cells; color, mean expression. (D) Schematic of the tier distance metric. Epithelial spots (green) anchor stepwise tier assignments to surrounding stromal spots (pink) on the hexagonal grid. (E) (Left) UMAP and (right) spatial map of stromal niche sub-clusters from spatially variable gene-based sub-clustering. (F) Partition-based graph abstraction (PAGA) of stromal sub-clusters. Edge thickness, connectivity. (G) UMAP colored by cell2location-inferred ECMi fraction. (H) Violin plots of ECMi (top) and ECMr (bottom) fractions across stromal sub-clusters. (I) UMAP colored by diffusion pseudotime. LISA, Local Indicators of Spatial Association; PAGA, partition-based graph abstraction.

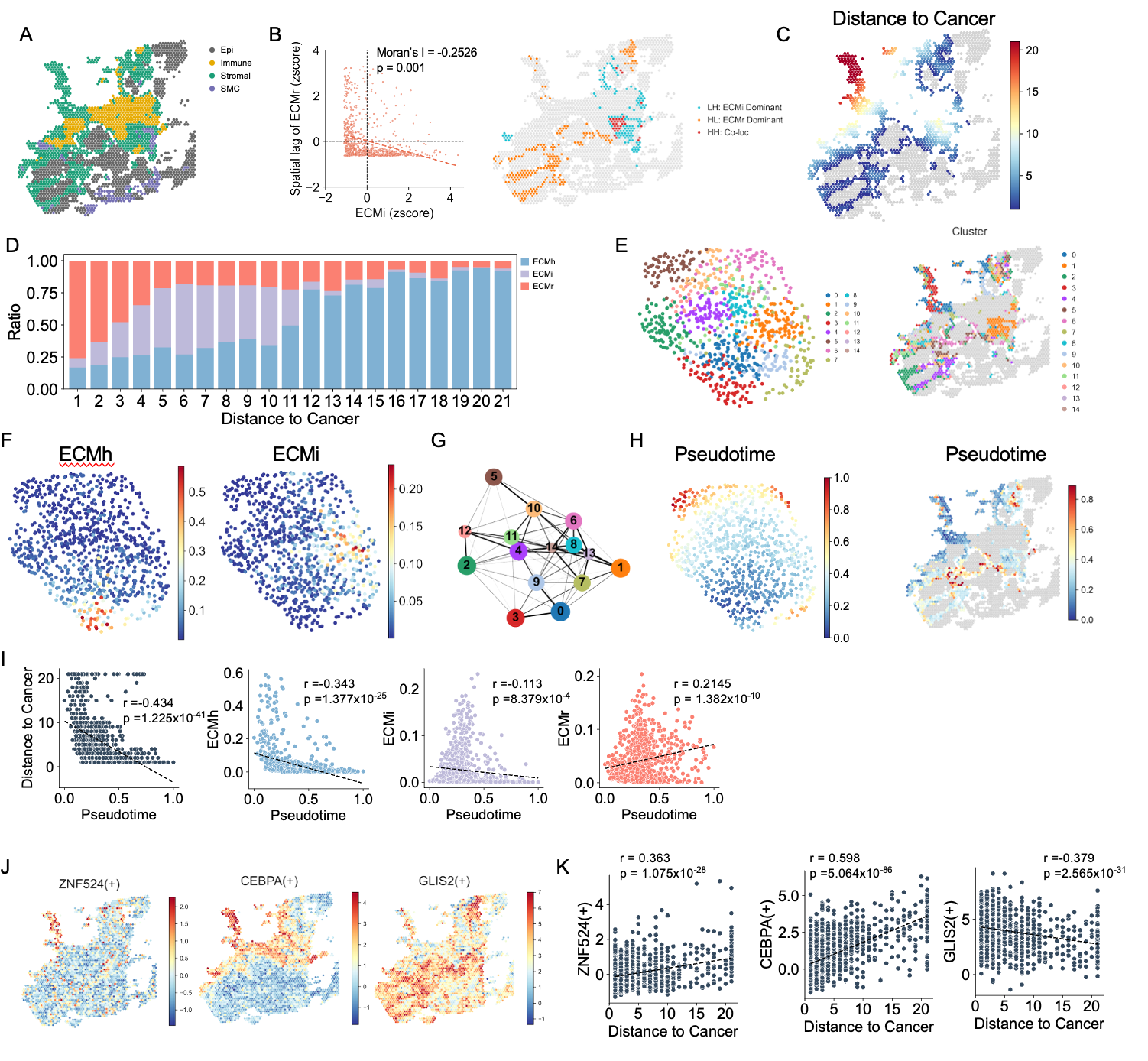

**Figure S6. ST2 spatial trajectory analysis.** (A) Spatial niche annotation in ST2 by LISA clustering. (B) (Left) Scatter plot of ECMi z-score against the spatial lag of ECMr z-score, with global bivariate Moran's I. (Right) Spatial map of spots classified by local bivariate Moran's I into ECMi-dominant, ECMr-dominant, and co-localized categories. (C) Distance to cancer map. (D) ECMh, ECMi, and ECMr ratio across distance to cancer bins in ST2 stromal niche spots. (E) (Left) UMAP and (right) spatial map of stromal niche sub-clusters. (F) UMAP colored by cell2location-inferred ECMh and ECMi fractions. (G) PAGA graph of stromal sub-clusters. Edge thickness, connectivity. (H) (Left) UMAP and (right) spatial map of diffusion pseudotime. (I) Scatter plots of distance to cancer, ECMh, ECMi, and ECMr fractions against pseudotime. Dashed lines, linear regression fit; Pearson r and P values are indicated. (J) Spatial maps of CEBPA(+) and GLIS2(+) regulon activity. (K) Scatter plots of CEBPA(+) and GLIS2(+) regulon activity against distance to cancer.

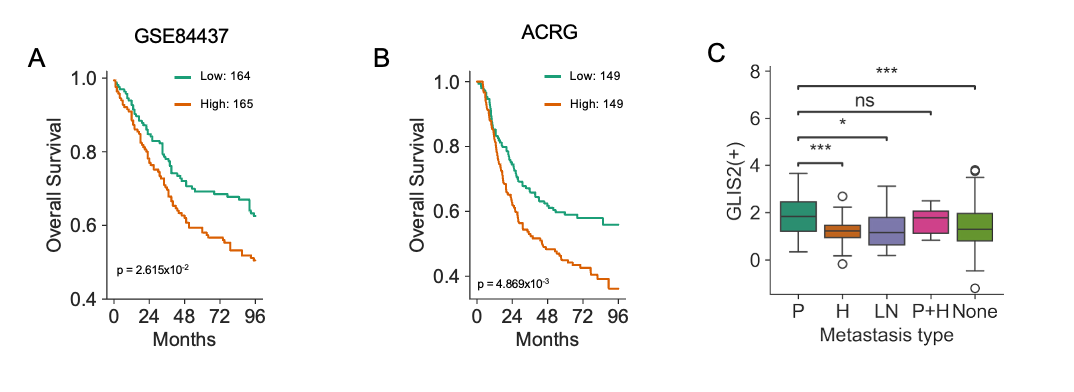

**Figure S7. Overall survival and metastasis type comparison by GLIS2(+) activity in gastric cancer cohorts.** (A) Kaplan-Meier curves of overall survival in the GSE84437 cohort, stratified into GLIS2(+) activity-high and -low groups by the median. Log-rank test. (B) As in (A) for the ACRG cohort. (C) GLIS2(+) regulon activity across metastasis types in the GSE84437 cohort. P, peritoneal; H, hematogenous; LN, lymph node; None, no recurrence. Center line, median; box, interquartile range; whiskers, 1.5 × interquartile range. Welch's t-test (*P < 0.05, ***P < 0.001; ns, not significant).

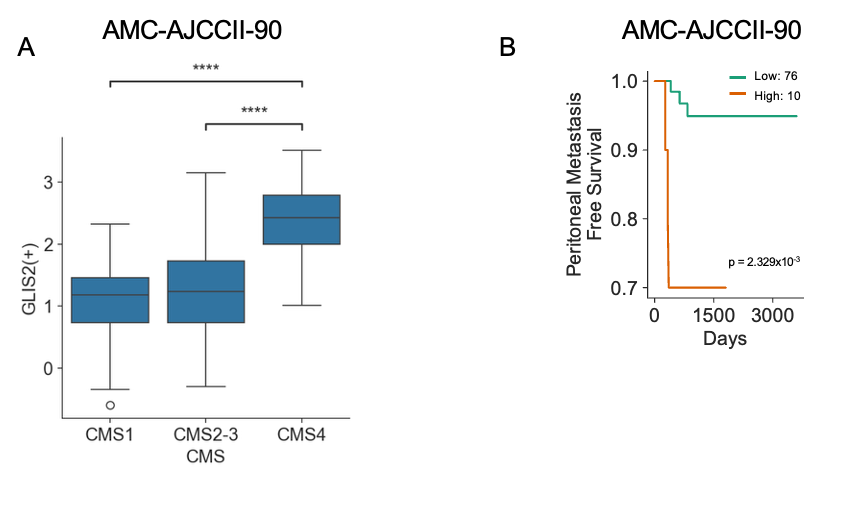

**Figure S8. GLIS2(+) activity in the colorectal cancer cohort AMC-AJCCII-90.** (A) GLIS2(+) regulon activity across consensus molecular subtypes (CMS1, CMS2-3, CMS4) of the AMC-AJCCII-90 cohort. Center line, median; box, interquartile range; whiskers, 1.5 × interquartile range. Welch's t-test (****P < 0.0001). (B) Kaplan-Meier curves of peritoneal metastasis-free survival in the AMC-AJCCII-90 cohort, stratified into GLIS2(+) activity-high and -low groups by the maximally selected rank statistic. Log-rank test.

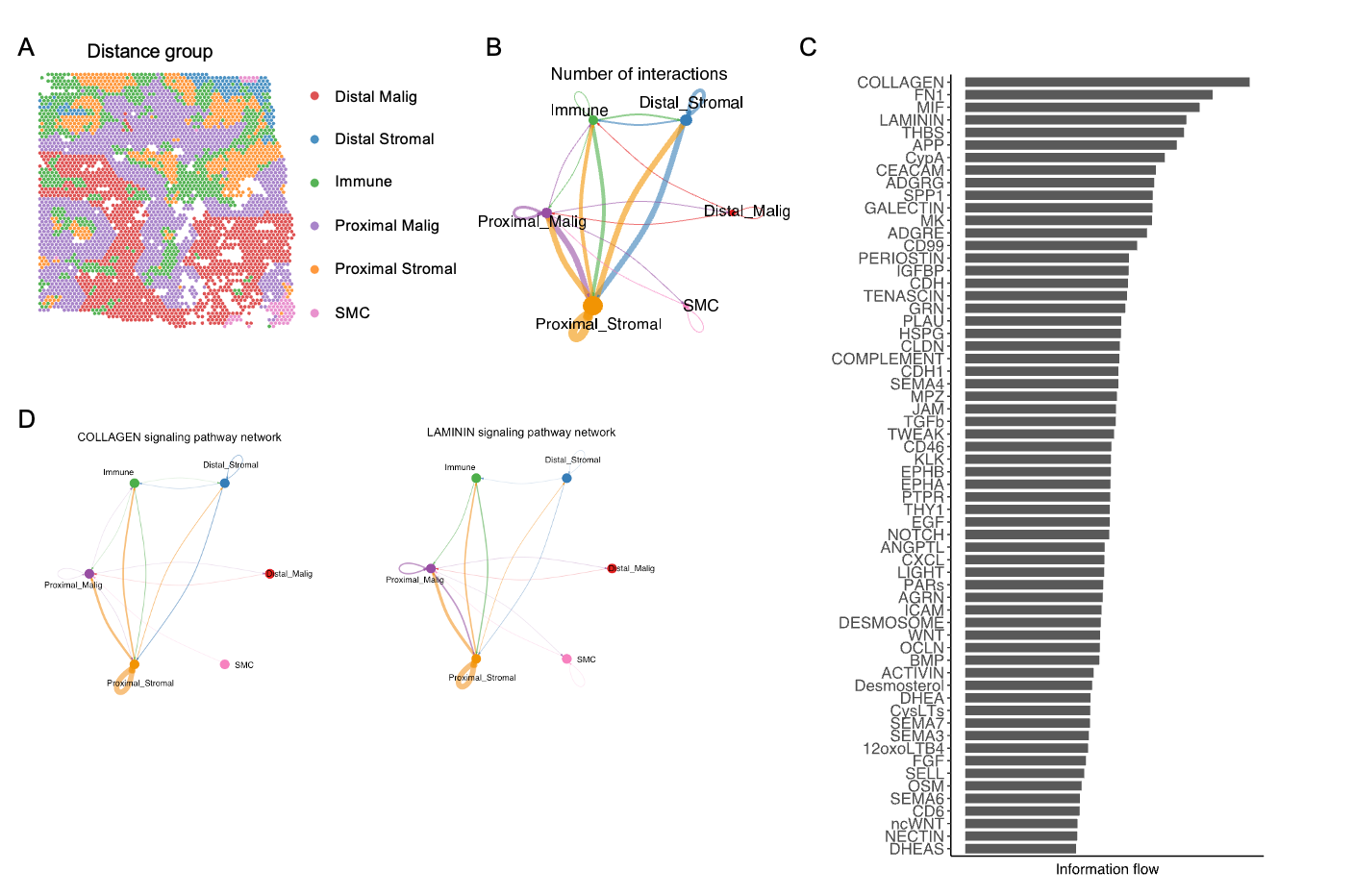

**Figure S9. Spatial cell-cell communication analysis in ST1.** (A) Spatial map of six niche groups in ST1 defined by proximity to epithelial spots (within five hexagonal steps as proximal). (B) Number of inferred interactions across the six niches. Edge thickness, interaction count. (C) Bar plot of signaling pathways ranked by information flow across niches. (D) Niche-niche network of COLLAGEN (left) and LAMININ (right) signaling pathways. Edge thickness, interaction strength.

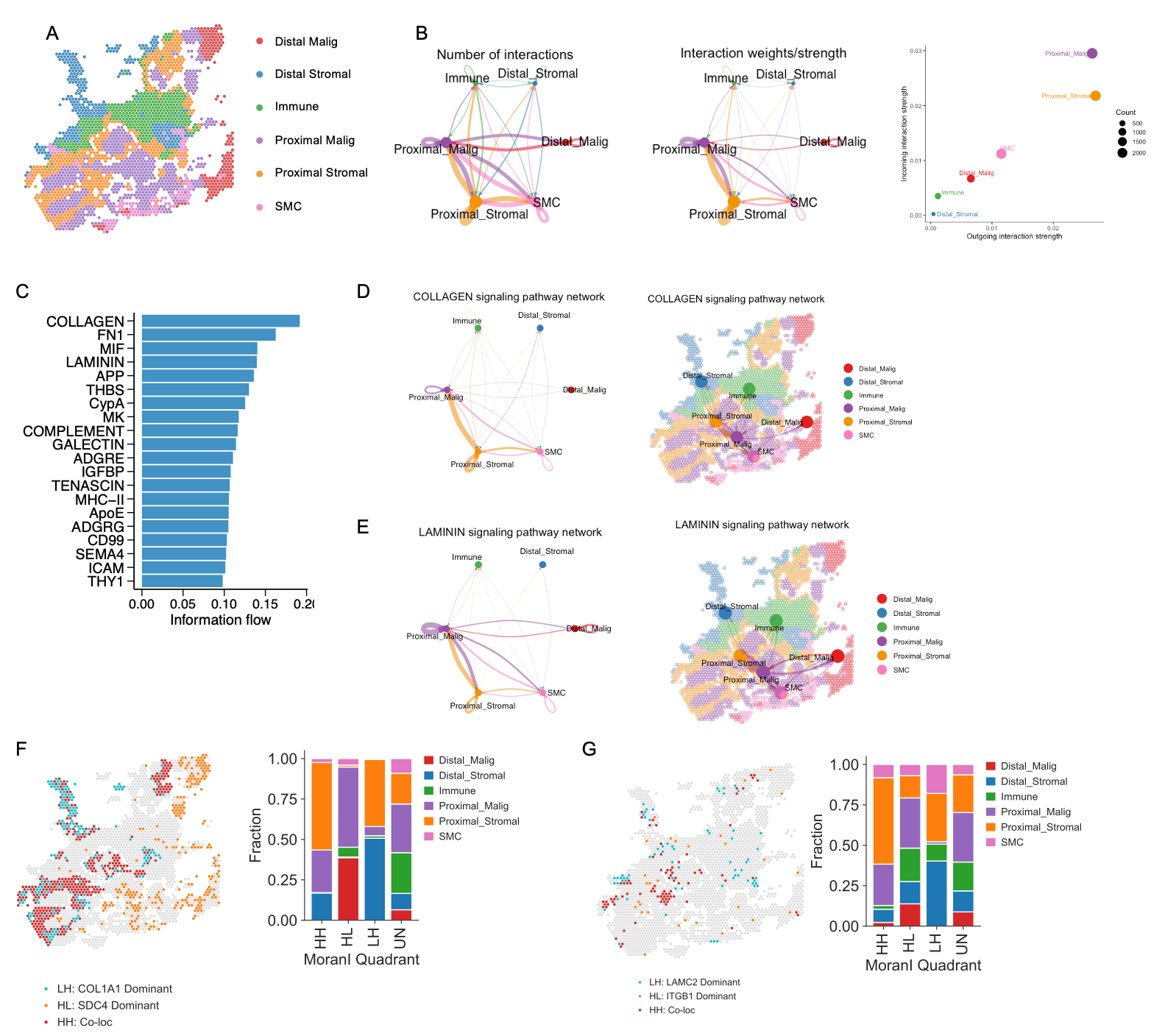

**Figure S10. Spatial cell-cell communication analysis in ST2.** (A) Spatial map of six niche groups in ST2. (B) (Left and middle) Niche-niche networks of number of interactions and interaction strength. (Right) Outgoing and incoming interaction strength across niches; dot size, number of inferred links. (C) Signaling pathways ranked by information flow across niches. (D) COLLAGEN signaling pathway network (left) and spatial map (right). (E) LAMININ signaling pathway network (left) and spatial map (right). (F) (Left) Local bivariate Moran's I spatial map of COL1A1 and SDC4 in ST2. (Right) Niche composition per Moran's I quadrant. (G) As in (F) for LAMC2 and ITGB1.

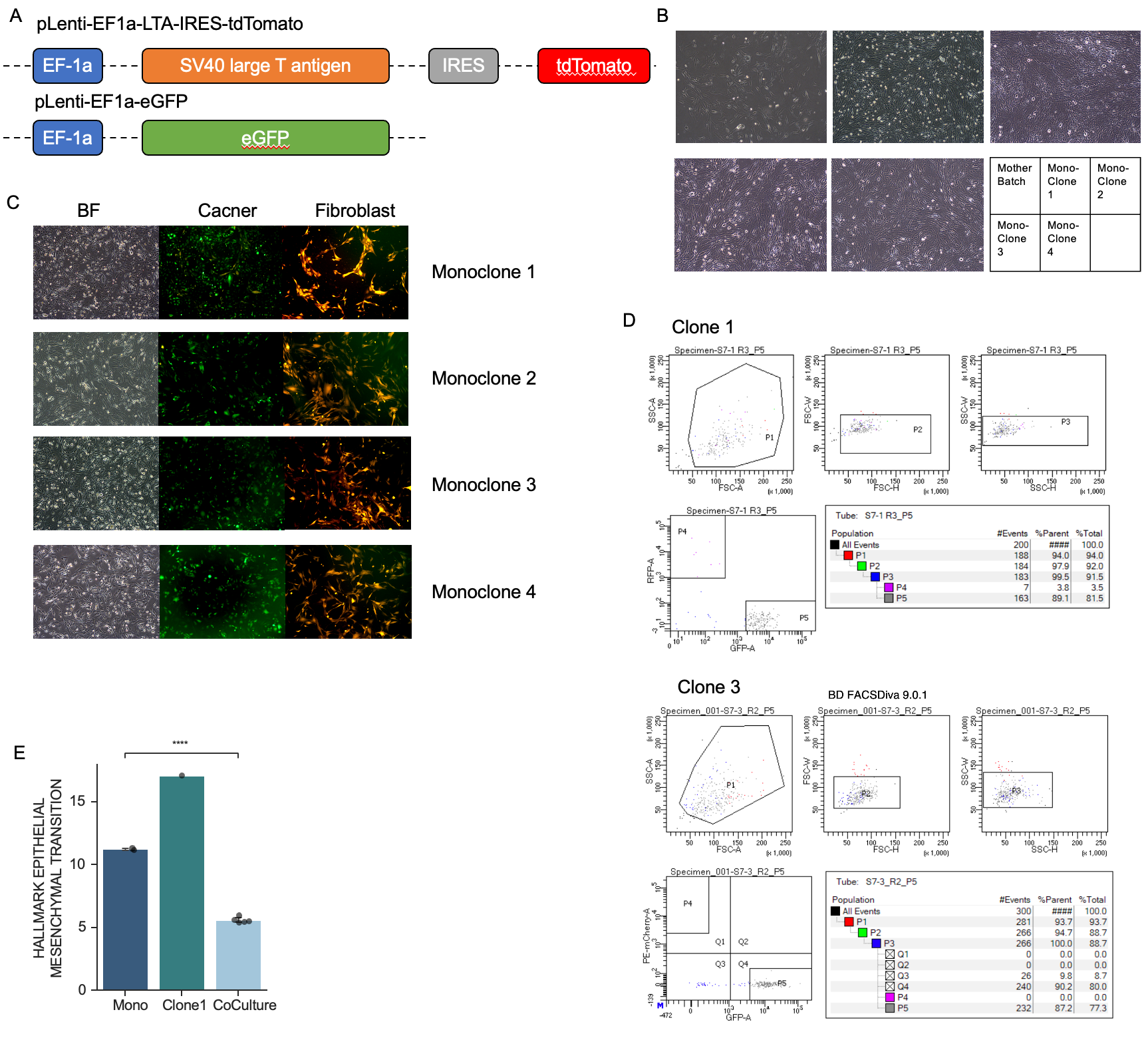

**Figure S11. Validation of patient-derived fibroblast lines and post-sort purity assessment.** (A) Vector schematics of pLenti-EF1a-LTA-IRES-tdTomato and pLenti-EF1a-eGFP. (B) Brightfield images of the parental immortalized pdFib pool and four monoclonal lines. (C) Brightfield and fluorescence images of SNU668 cancer cells (GFP) co-cultured with monoclonal pdFib lines (tdTomato). (D) Post-sort flow cytometry gating of GFP+ SNU668 cells after co-culture with clone 1 (top) and clone 3 (bottom), assessing residual fibroblast contamination. (E) Hallmark epithelial-mesenchymal transition score in monoculture (Mono), clone 1 co-cultured and clone 2 to 4 co-cultured (CoCulture) SNU668 cells. Error bars, 95% confidence intervals. Welch's t-test (****P < 0.0001).

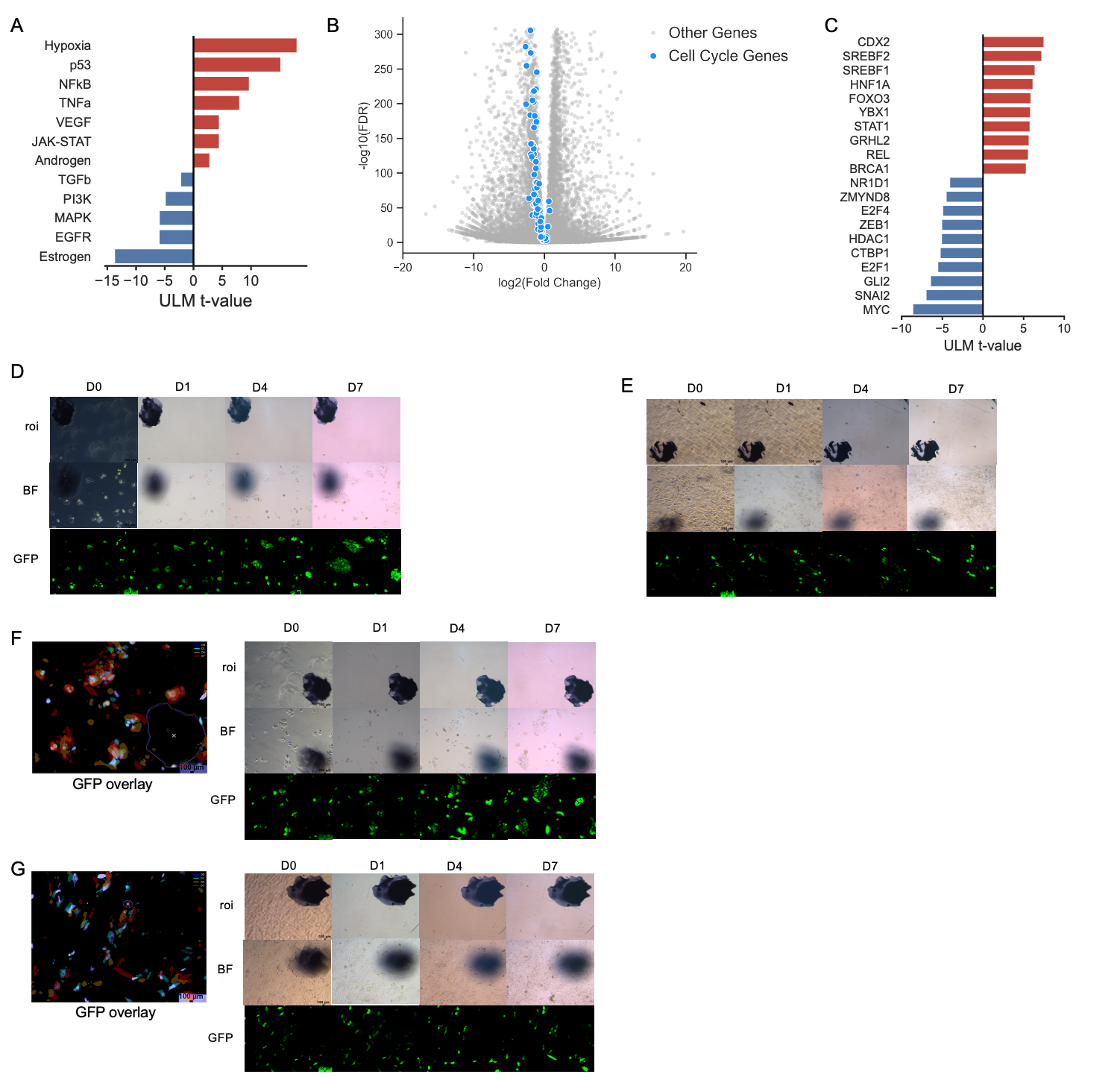

**Figure S12. Additional pathway, transcription factor, and longitudinal imaging analyses of co-cultured SNU668 cells.** (A) PROGENy signaling pathway activity in co-cultured versus WT SNU668 cells, ranked by ULM t-value. (B) Volcano plot of differential gene expression in co-cultured versus WT SNU668 cells. Cell cycle genes are highlighted (blue). (C) Top up- and down-regulated CollecTRI transcription factor activity in co-cultured versus WT SNU668 cells, ranked by ULM t-value. (D and E) Longitudinal brightfield (BF) and GFP imaging of SNU668 cells in monoculture at D0, D1, D4, and D7 in two independent replicate regions. Scale bar, 100 µm. (F and G) Longitudinal imaging of SNU668 cells co-cultured with ECMr fibroblasts in two independent replicate regions. (Left) GFP overlay across D0, D1, D4, and D7, with each time point assigned a distinct color. (Right) BF and GFP channels at each time point. Scale bar, 100 µm.

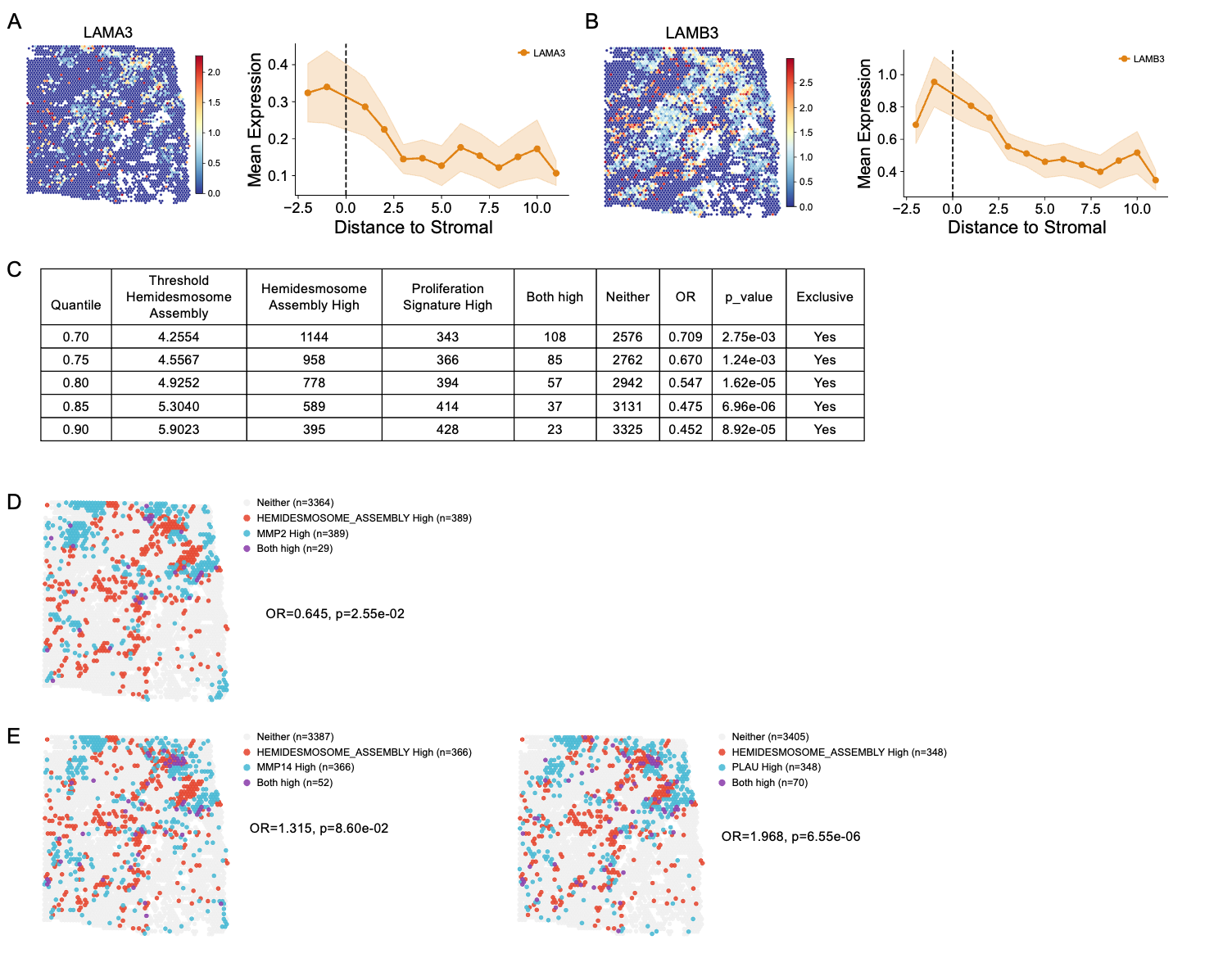

**Figure S13. Spatial validation of LM332 components and wound healing-type protease pattern in ST1.** (A) (Left) Spatial map of LAMA3 expression in ST1. (Right) Mean LAMA3 expression across distance to stromal. Shaded area, 95% confidence interval. (B) As in (A) for LAMB3. (C) Sensitivity analysis of the inverse relationship between hemidesmosome assembly and proliferation signature scores across hemidesmosome assembly quantile thresholds. Proliferation signature threshold fixed at zero. Fisher's exact test. (D) Spatial map of epithelial niche spots classified by hemidesmosome assembly and MMP2 expression. Fisher's exact test. (E) As in (D) for MMP14 (left) and PLAU (right).

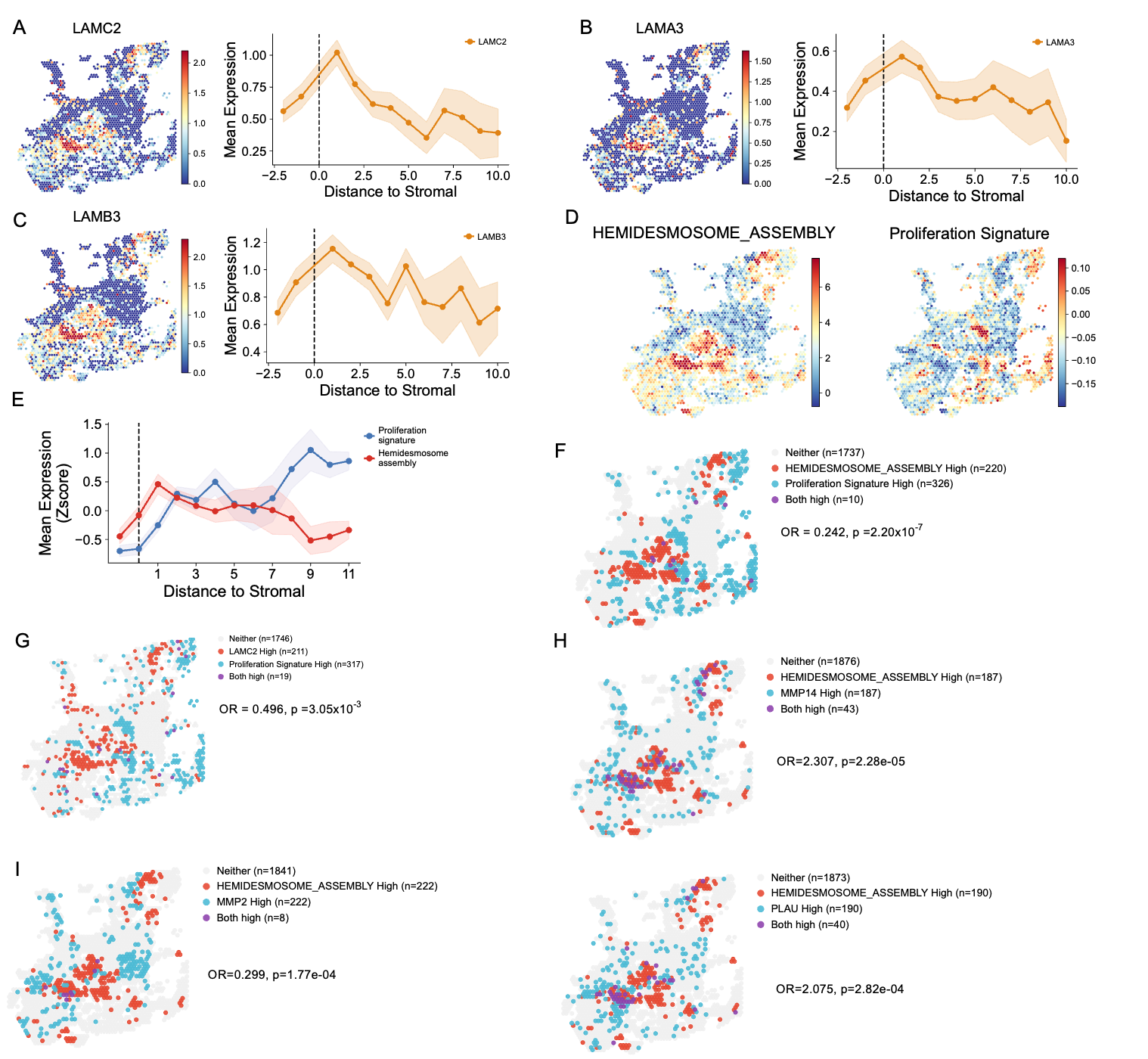

**Figure S14. Reproduction of LM332, hemidesmosome assembly, and protease spatial patterns in ST2.** (A) (Left) Spatial map of LAMC2 expression in ST2. (Right) Mean LAMC2 expression across distance to stromal. Shaded area, 95% confidence interval. (B and C) As in (A) for LAMA3 and LAMB3. (D) Spatial maps of hemidesmosome assembly (left) and proliferation signature (right) scores in ST2. (E) Mean hemidesmosome assembly and proliferation signature scores (z-scored) across distance to stromal. Shaded areas, 95% confidence intervals. (F) Spatial map of epithelial niche spots classified by hemidesmosome assembly (90th percentile threshold) and proliferation signature (zero threshold). Fisher's exact test. (G) As in (F) for LAMC2 and proliferation signature. (H) As in (F) for hemidesmosome assembly and MMP14. (I) As in (F) for hemidesmosome assembly and MMP2 (left) and PLAU (right).

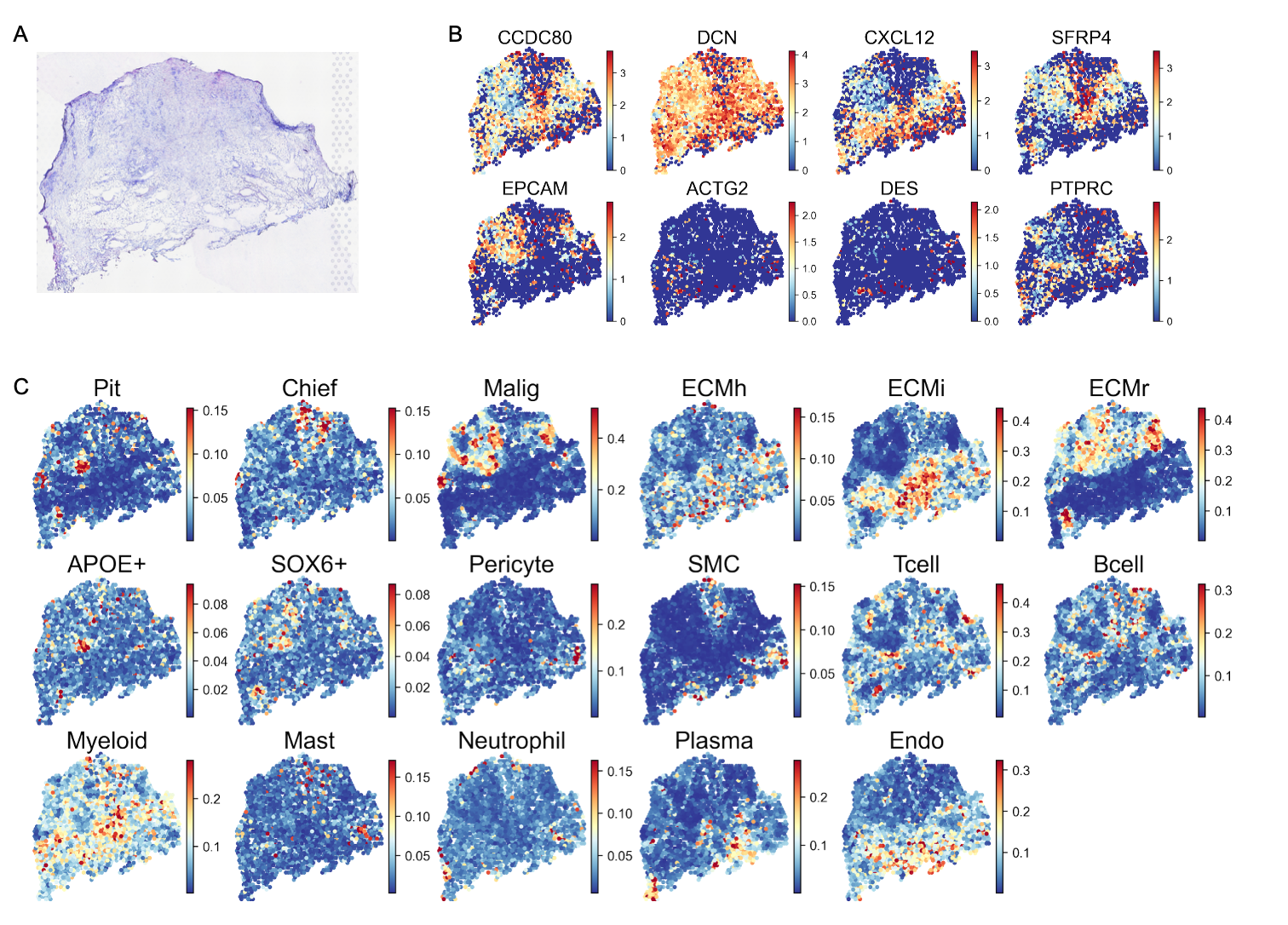

**Figure S15. Histology and spatial cell type maps of PT1.** (A) H&E image of PT1. (B) Spatial expression maps of stromal (CCDC80, DCN, CXCL12, SFRP4), epithelial (EPCAM), smooth muscle (ACTG2, DES), and immune (PTPRC) marker genes. Color, log-normalized expression. (C) Cell2location-inferred spatial fraction maps of epithelial subsets (Pit, Chief, Malig), ECM fibroblasts (ECMh, ECMi, ECMr), other stromal subsets (APOE+, SOX6+, Pericyte, SMC), immune subsets (Tcell, Bcell, Myeloid, Mast, Neutrophil, Plasma), and endothelial cells (Endo). Color, estimated cell fraction per spot.

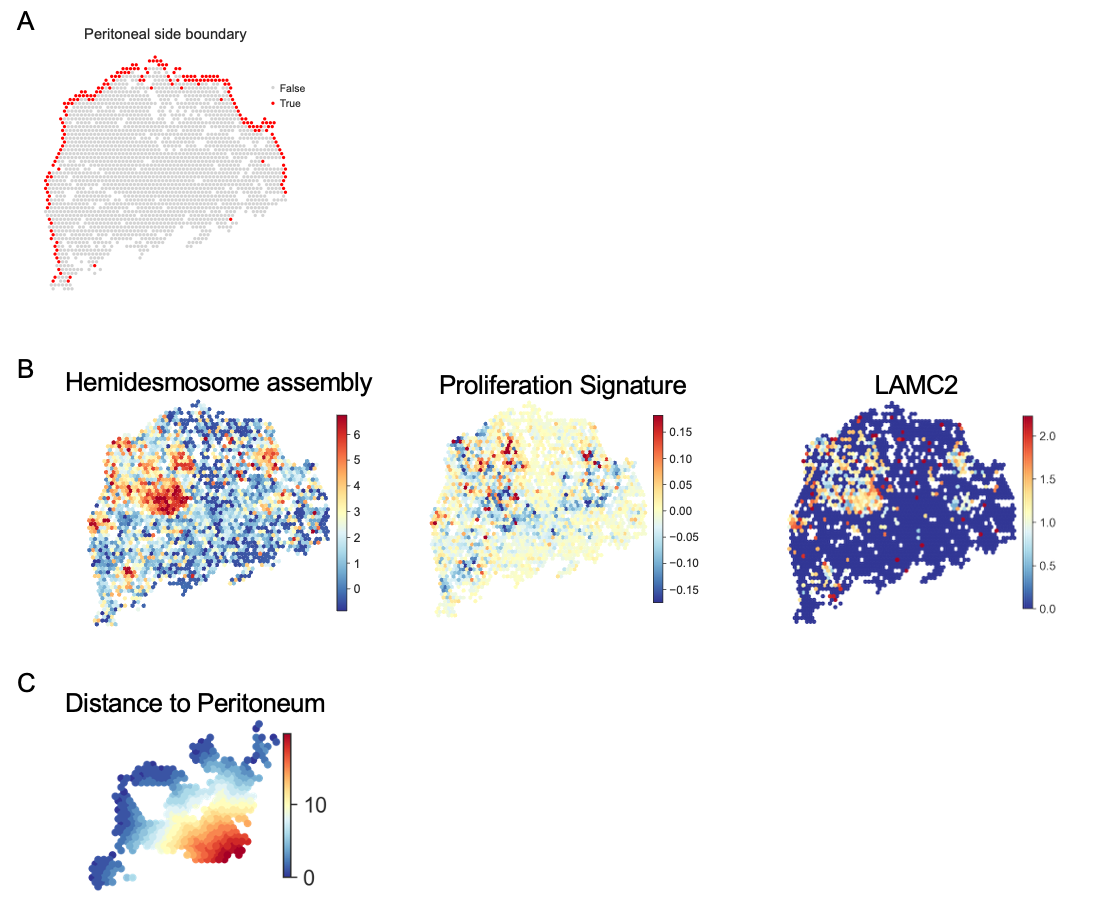
**Figure S16. Spatial validation of hemidesmosome assembly, proliferation, and LAMC2 in PT1.** (A) Spatial map of PT1 with peritoneal side boundary spots highlighted (red). (B) Spatial maps of hemidesmosome assembly score, proliferation signature score, and LAMC2 expression within the epithelial niche region of interest in PT1. (C) Spatial map of distance to peritoneal surface within the same region of interest as in Fig. 8G.

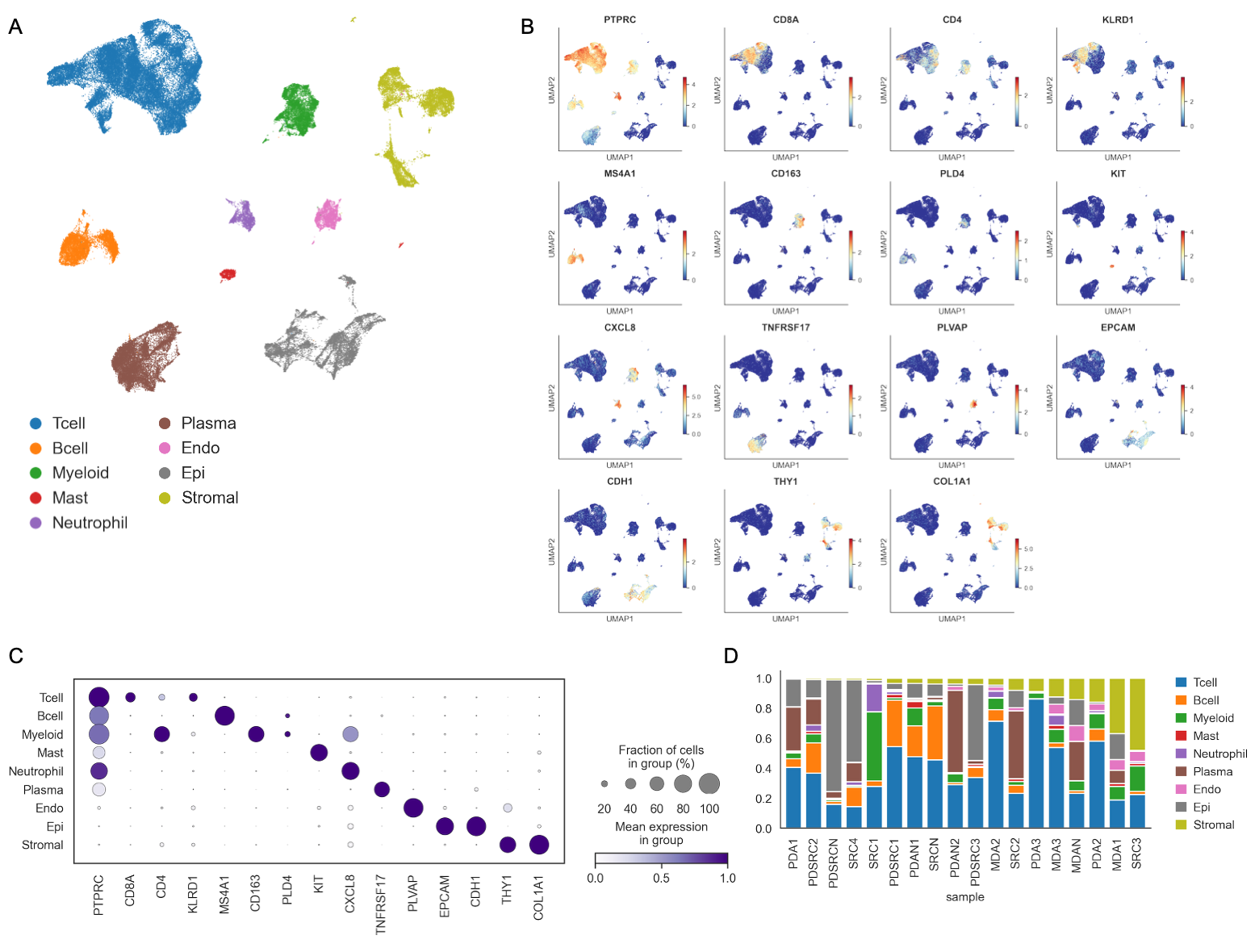

**Figure S17. Major cell type annotation in gastric cancer scRNA-seq data.** (A) UMAP of all cells, colored by major cell type. (B) UMAP feature plots of canonical marker genes. Color, log-normalized expression. (C) Dot plot of marker gene expression across major cell types. Dot size, fraction of cells; color, scaled mean expression. (D) Stacked bar plot of major cell type composition per patient sample.

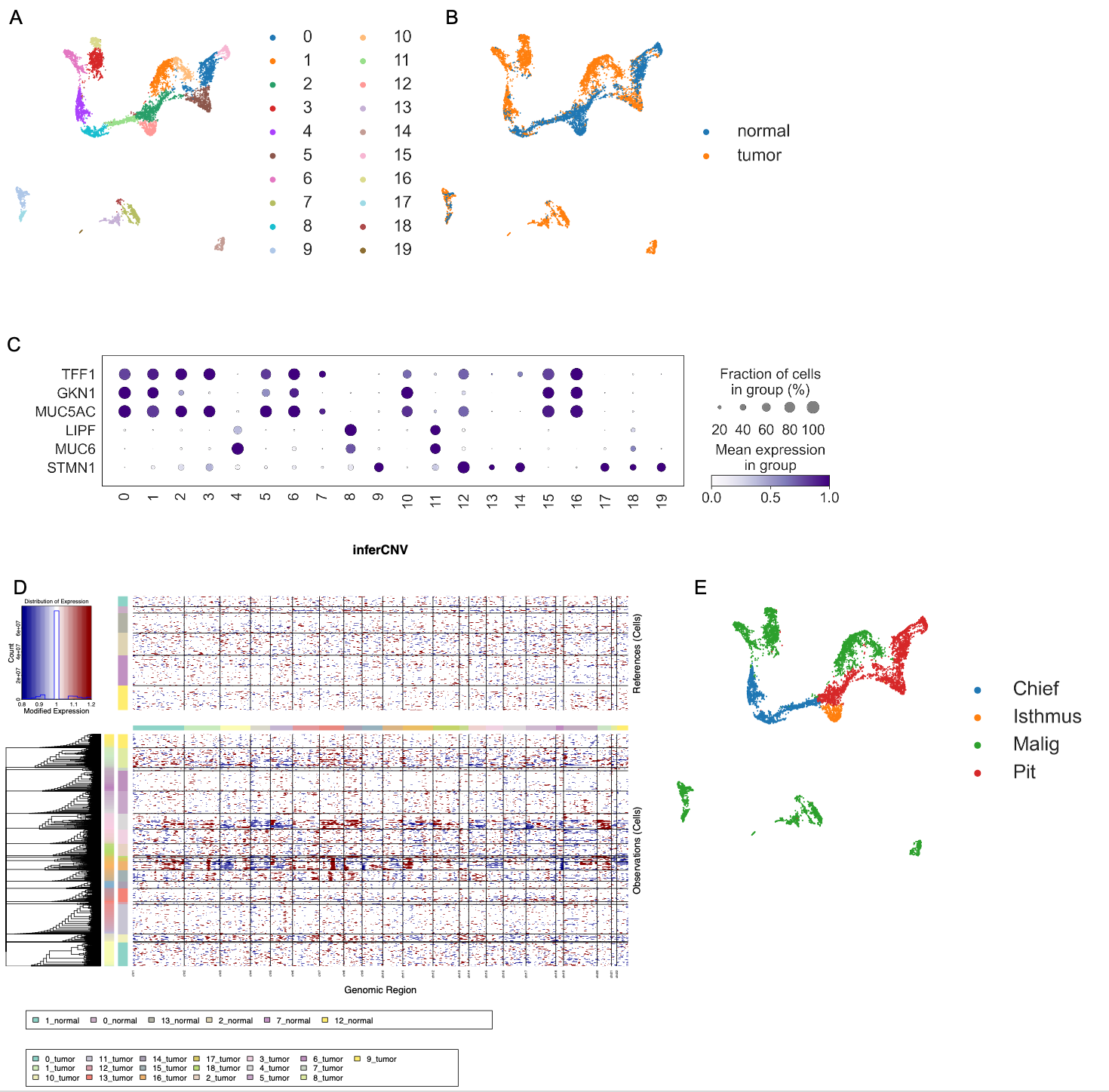

**Figure S18. Epithelial cell clustering and inferCNV-based malignant cell identification.** (A) UMAP of epithelial cells, colored by Leiden sub-cluster. (B) UMAP colored by sample origin (normal or tumor). (C) Dot plot of canonical gastric epithelial marker gene expression across sub-clusters. TFF1, GKN1, MUC5AC, pit cell; LIPF, MUC6, chief and gland mucous cell; STMN1, isthmus and proliferative compartment. Dot size, fraction of cells; color, scaled mean expression. (D) inferCNV heatmap of copy number alteration across genomic regions. Top, reference cells from normal-origin clusters. Bottom, observation cells from tumor-origin clusters. Side bar indicates sub-cluster identity. (E) UMAP colored by epithelial cell type annotation (Chief, Isthmus, Malig, Pit).

**Supplementary Tables**

**Table S1. Clinical characteristics and bulk gene expression of the spatial transcriptomics sample cohort.**

Clinicopathologic features (sex, age, AGC type, histologic type, Lauren classification, T/N/M stage, TNM stage) and gene expression values of SFRP4, GZMB, WARS, and CDX1 for samples ST1, ST2, PT1, and NT1. Each row corresponds to one tissue sample.

**Table S2. Marker genes of stromal cell subsets identified by single-cell RNA-seq.**

Differentially expressed genes for each stromal subset from Seurat FindAllMarkers analysis. Columns: average log2 fold change (avg_log2FC), fraction of expressing cells in the cluster (pct.1) and in all other clusters (pct.2), unadjusted P value (p_val), Bonferroni-adjusted P value (p_val_adj), cluster identity, and gene symbol.

**Table S3. Differential pathway activity between ECMi and ECMr fibroblasts**.

GSEA-derived pathway-level t-statistics and P values for the comparison of ECMi versus ECMr fibroblast subsets. Positive t-statistics indicate higher activity in ECMr; negative values indicate higher activity in ECMi.

**Table S4. CytoTRACE2 stemness scores and diffusion pseudotime in stromal cells.**

CytoTRACE2 outputs (CytoTRACE2_Score, CytoTRACE2_Potency, CytoTRACE2_Relative, and corresponding preKNN values) and diffusion pseudotime values (dpt_pseudotime) computed for each stromal cell. Each row corresponds to one cell.

**Table S5. SCENIC-derived gene regulatory network.**

Inferred transcription factor-to-target gene relationships from SCENIC analysis. Columns: regulator transcription factor (source), target gene (target), and regulatory weight (weight).

**Table S6. BEAM analysis of branch-dependent regulon activity.**

Branched expression analysis modeling (BEAM) results testing which regulons diverge between the two branches of the stromal trajectory. Columns: regulon name, mean absolute error of divergence (mae_divergence) and its normalized form, pooled standard deviation, permutation P value, number of valid permutations, number of cells assigned to branch 1 and branch 2, and Benjamini-Hochberg adjusted P value (qvalue).

**Table S7. CellChat ligand-receptor interactions across cell populations.**

Inferred ligand-receptor pairs from CellChat analysis. Columns: source cell population, target cell population, ligand, receptor, interaction probability (prob), permutation P value, interaction name (raw and formatted), pathway name, annotation category, and supporting evidence.

**Table S8. Differential gene expression in ECMr-co-cultured versus monocultured SNU668 cells.**

pyDESeq2 results comparing SNU668 cells co-cultured with ECMr fibroblasts against monocultured SNU668 cells. Columns: gene symbol, baseMean expression, log2 fold change, log2 fold change standard error (lfcSE), Wald statistic (stat), unadjusted P value, and Benjamini-Hochberg adjusted P value (padj).

**Table S9. Oligonucleotide and primer sequences.**

Oligonucleotides and primers used for cloninglentiviral construct generation. Each row lists the name and the 5′ to 3′ sequence.
